# Defective glycosylation and ELFN1 binding of mGluR6 congenital stationary night blindness mutants

**DOI:** 10.1101/2024.10.28.620767

**Authors:** Mustansir Pindwarawala, Faiyaz A. K. Abid, Jaeeun Lee, Michael L. Miller, Juliet S. Noppers, Andrew P. Rideout, Melina A. Agosto

## Abstract

Synaptic transmission from photoreceptors to ON-bipolar cells (BCs) requires the postsynaptic metabotropic glutamate receptor mGluR6, located at BC dendritic tips. Binding of the neurotransmitter glutamate initiates G-protein signaling that regulates the TRPM1 transduction channel. mGluR6 also interacts with presynaptic ELFN adhesion proteins, and these interactions are important for mGluR6 synaptic localization. The mechanisms of mGluR6 trafficking and synaptic targeting remain poorly understood. In this study, we investigated mGluR6 missense mutations from patients with congenital stationary night blindness (CSNB), which is associated with loss of synaptic transmission to ON-BCs. We found that multiple CSNB mutations in the extracellular ligand-binding domain of mGluR6 impart a trafficking defect leading to lack of complex N-glycosylation but efficient plasma membrane insertion, suggesting a Golgi bypass mechanism. These mutants fail to bind ELFN1, consistent with lack of a necessary modification normally acquired in the Golgi. The same mutants were mislocalized in bipolar cells, explaining the loss of function in CSNB. The results reveal a key role of Golgi trafficking in mGluR6 function, and suggest a role for the extracellular domain in Golgi sorting.

## Introduction

In the retina outer plexiform layer (OPL), signals from rod and cone photoreceptors are transmitted to bipolar cells through synaptic release of glutamate. The detection of glutamate at the dendritic tips of ON-type bipolar cells (ON-BCs) is mediated by metabotropic glutamate receptor 6 (mGluR6) (1–4), a class C G-protein coupled receptor. mGluR6 is coupled to G_o_ (5–7), and upon agonism by glutamate in the dark, G_αo_ and/or G_βγ_ inhibit TRPM1 channel opening via an unknown mechanism (8–13). Deactivation of mGluR6 in the light relieves the inhibition, leading to ON-BC depolarization at light onset. In addition to its critical role in neurotransmitter detection, mGluR6 binds in trans with presynaptic ELFN1 and ELFN2, and these interactions are important for synapse formation and mGluR6 synaptic enrichment (14, 15).

The mGluR6 protein has a large extracellular domain consisting of a bilobed ligand binding domain (LBD) and a membrane-proximal cysteine-rich domain (CRD), and like all GPCRs, a 7-pass transmembrane domain (TMD) (16). Previous work from our lab demonstrated that ELFN1 and ELFN2 preferentially interact with mGluR6 that has been modified by complex N-linked glycosylation in the Golgi (17). We found that four sites in the extracellular domain are N-glycosylated; the glycan at N445 (N451 in human mGluR6) was of particular importance for ELFN1 and ELFN2 binding, and was sufficient for partial synaptic enrichment in BCs (17).

The majority of transmembrane proteins destined for the plasma membrane (PM) are modified by N-glycosylation, acquired first in the endoplasmic reticulum (ER), then modified to complex forms in the Golgi (18). However, some proteins follow an unconventional protein secretion (UPS) pathway that bypasses the Golgi, either as part of their normal trafficking, or due to ER stress (19, 20). The mechanisms of sorting to the Golgi versus a UPS pathway are poorly understood.

Congenital stationary night blindness (CSNB) is a condition characterized by impaired synaptic transmission from photoreceptors to ON-BCs (21). Numerous alleles of the *GRM6* gene, which codes for mGluR6, have been associated with autosomal recessive complete CSNB (22–31). CSNB cases with *GRM6* mutations are characterized by an electronegative waveform (no b-wave) in electroretinograms, indicating loss of ON-BC postsynaptic function. However, the molecular basis for this loss of function is largely unknown. Mutations in genes encoding other ON-BC synaptic proteins, including TRPM1, nyctalopin, GPR179, and LRIT3, are also associated with complete CSNB (21). While ON-BCs participate in both the rod and cone pathways, cone neurotransmission to OFF type BCs is spared in complete CSNB. The rod pathway, in contrast, depends on rod ON-BCs, leading to selective dysfunction in dim light.

In this study, we characterized a panel of 16 missense CSNB mutations in mGluR6. We found that most of the mutations in the mGluR6 ligand-binding domain are competent for trafficking to the plasma membrane, but have defective N-linked glycosylation, fail to bind to ELFN1, and are mislocalized in ON-BCs.

## Results

### Most LBD mutants are competent for PM trafficking in HEK cells but are mislocalized in ON-BCs

A panel of 16 missense CSNB mutants (22–29) of human mGluR6 was constructed by site-directed mutagenesis. To test PM trafficking, HEK cells transfected with untagged human mGluR6 were labeled in non-permeabilizing conditions with mAb-1438, which recognizes an extracellular epitope in both mouse and human mGluR6 (32) (Fig S1). In contrast to a previous report (24), we observed robust surface expression of many of the variants with mutations in the LBD (Fig 1). With the exception of G275D and C423R, all of the LBD mutants had at least ∼35% of normal plasma membrane protein, and that of two mutants – G150S and R352C – was similar to or better than WT. The remaining mutations, located in the CRD or TMD, significantly reduced surface expression. Total expression, assayed by labeling in permeabilizing conditions, was not greatly affected for any of mutants (Fig 1). Surface expression of mGluR6-EGFP fusion proteins was similar to the corresponding untagged mutants, though the CRD and TMD mutations also reduced total expression measured by EGFP fluorescence (Fig S2). Like other mGluRs, mGluR6 likely forms dimers. However, untagged LBD mutants, regardless of whether they had superior or defective surface expression, had no apparent dominant effect on PM levels of co-transfected WT Myc-tagged mGluR6 in HEK cells (Fig S3).

**Figure 1.**
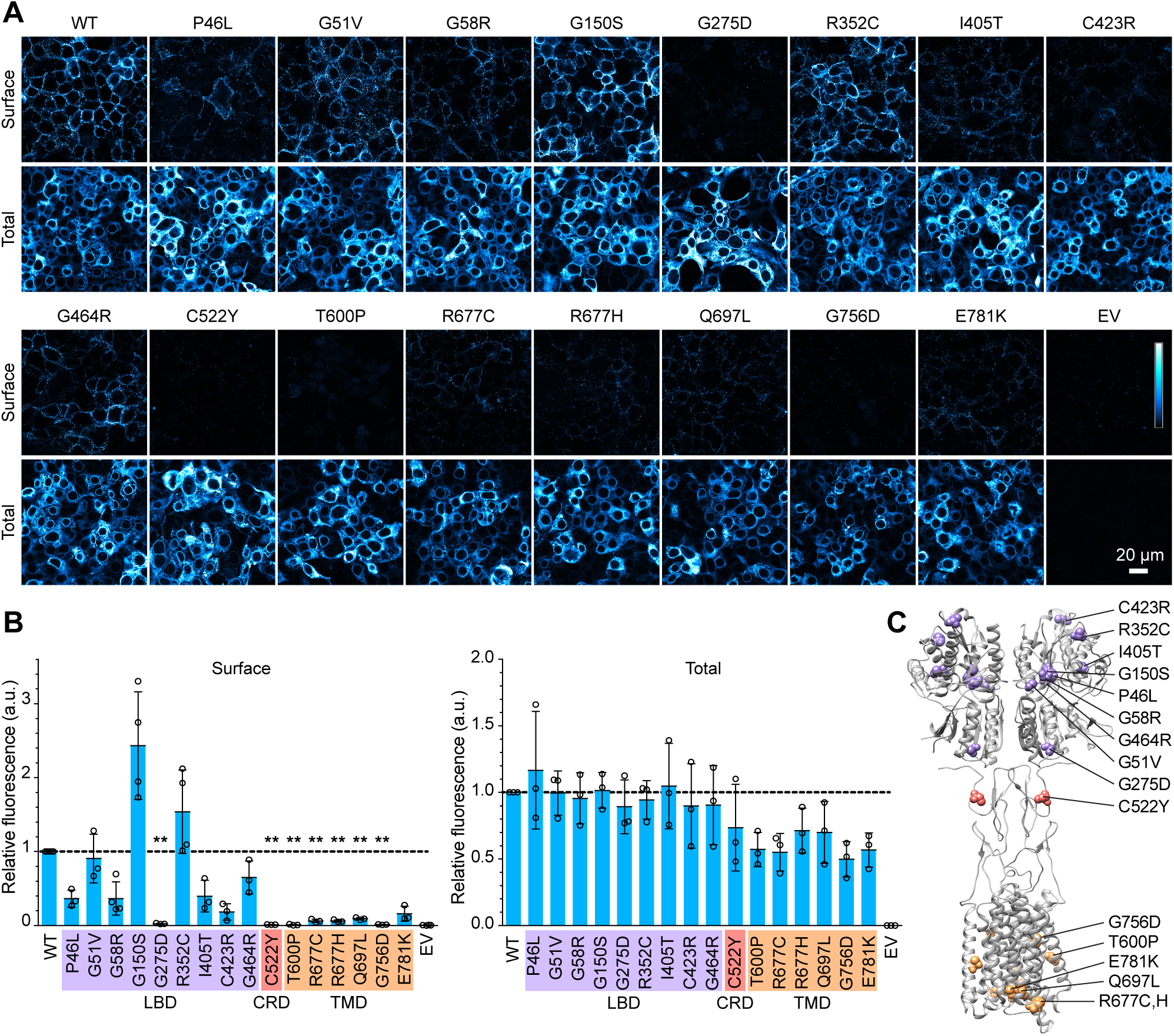
Surface expression of CSNB mutants in HEK cells. **(A)** Example images of cells transiently transfected with WT or mutant untagged mGluR6, or empty vector control (EV), and labeled with mGluR6 mAb 1438 in non-permeabilizing or permeabilizing conditions, to detect surface and total protein, respectively. **(B)** Quantification of images as shown in (A). Each point represents the mean of four images from an independent experiment, normalized to the WT samples from the same experiment. Mutants were compared to the WT value of 1 using 1-sample t-tests, and p-values were multiplied by 16 to correct for multiple comparisons. **, p<0.01. **(C)** Diagram of mGluR4 (PDB 7E9H (55)) with homologous positions of CSNB mutations shown as red spheres.

Next, we tested localization in BCs by in vivo electroporation of mGluR6-EGFP expression constructs in mouse retina (Fig 2). WT human mGluR6-EGFP was largely colocalized with TRPM1 in ON-BC dendritic tips, as we reported previously for electroporated murine mGluR6-EGFP (17, 32, 33). The subset of LBD mutants that we tested, which all had good to excellent surface expression in HEK cells, were severely mislocalized in BCs in both WT mice (Fig 2A,F) and mGluR6 null mice (Fig S4). TRPM1 dendritic tip localization, as well as total EGFP in the imaged fields, were similar between WT and mutants (Fig 2D,E). These results are consistent with our previous findings that competence for PM trafficking is not sufficient for synaptic localization of mGluR6 in BCs (32).

**Figure 2.**
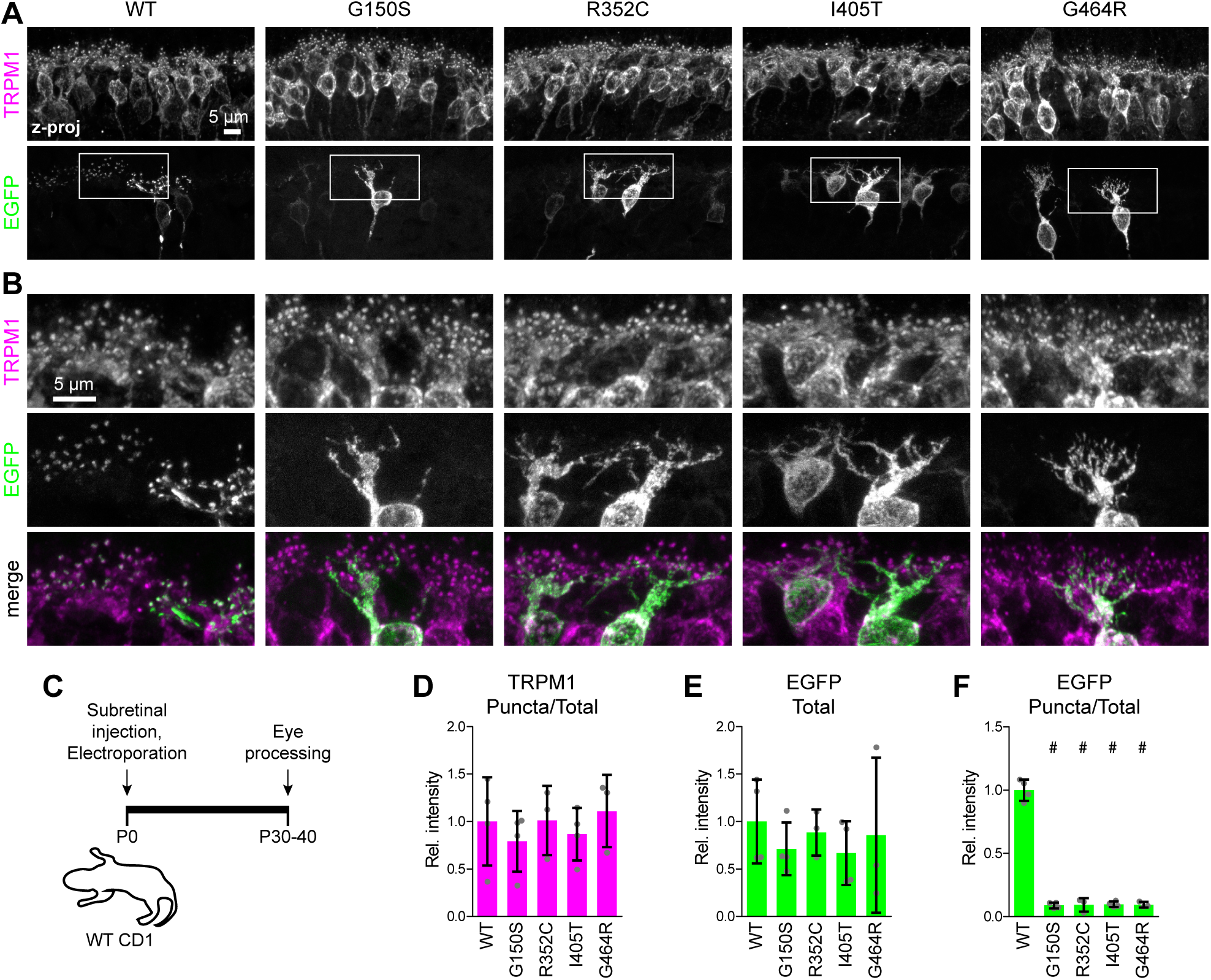
LBD mutants are mislocalized in mouse retinal BCs. **(A)** Example images of WT CD1 retinas electroporated with WT or mutant mGluR6-EGFP and labeled with TRPM1 antibody. **(B)** Magnified views of boxed regions in (A). **(C)** Diagram of electroporation procedure. **(D-F)** Quantification of images as shown in (A). Data are shown relative to the mean of WT mGluR6-EGFP. Each point represents the mean of at least three images from a different animal (n=3-4 biological replicates per condition), and bars represent means of biological replicates ± S.D. Mutants were compared to WT by one-way ANOVA and Dunnett’s post-test; #, p<0.001.

### LBD mutants have defects in N-glycosylation and ELFN1 binding

The mechanisms of mGluR6 targeting to BC postsynapses are unknown. However, defective binding to presynaptic ELFN proteins, or absence of ELFN, is associated with mGluR6 mislocalization (14, 15, 17, 32). Previously, in experiments done with murine proteins expressed in HEK cells, we reported that N-linked core-glycosylated (band B) and complex-glycosylated (band C) forms of WT mGluR6 are resolved by SDS-PAGE, and that ELFN1 specifically interacts with band C (17). Similarly, the extracellular domain of both human and mouse ELFN1 pulled down the complex glycosylated form (band C) of human mGluR6, but not band B (Fig 3A). When we tested CSNB mutants for binding to human ELFN1, we observed that band C appeared to be absent in the LBD mutants (Fig 3B, input samples). Consistent with selective interaction with band C, none of the LBD mutants had detectable ELFN1 binding (Fig 3B,C). We also tested three TMD mutants (R677C, Q697L, and E781K), which had significantly reduced surface expression in HEK cells (see Fig 1). In contrast to the LBD mutants, these TMD mutants did have an observable band C, and also exhibited some ELFN1 binding (Fig 3C,D). No nonspecific binding to negative control Fc protein was observed for any of the mutants.

**Figure 3.**
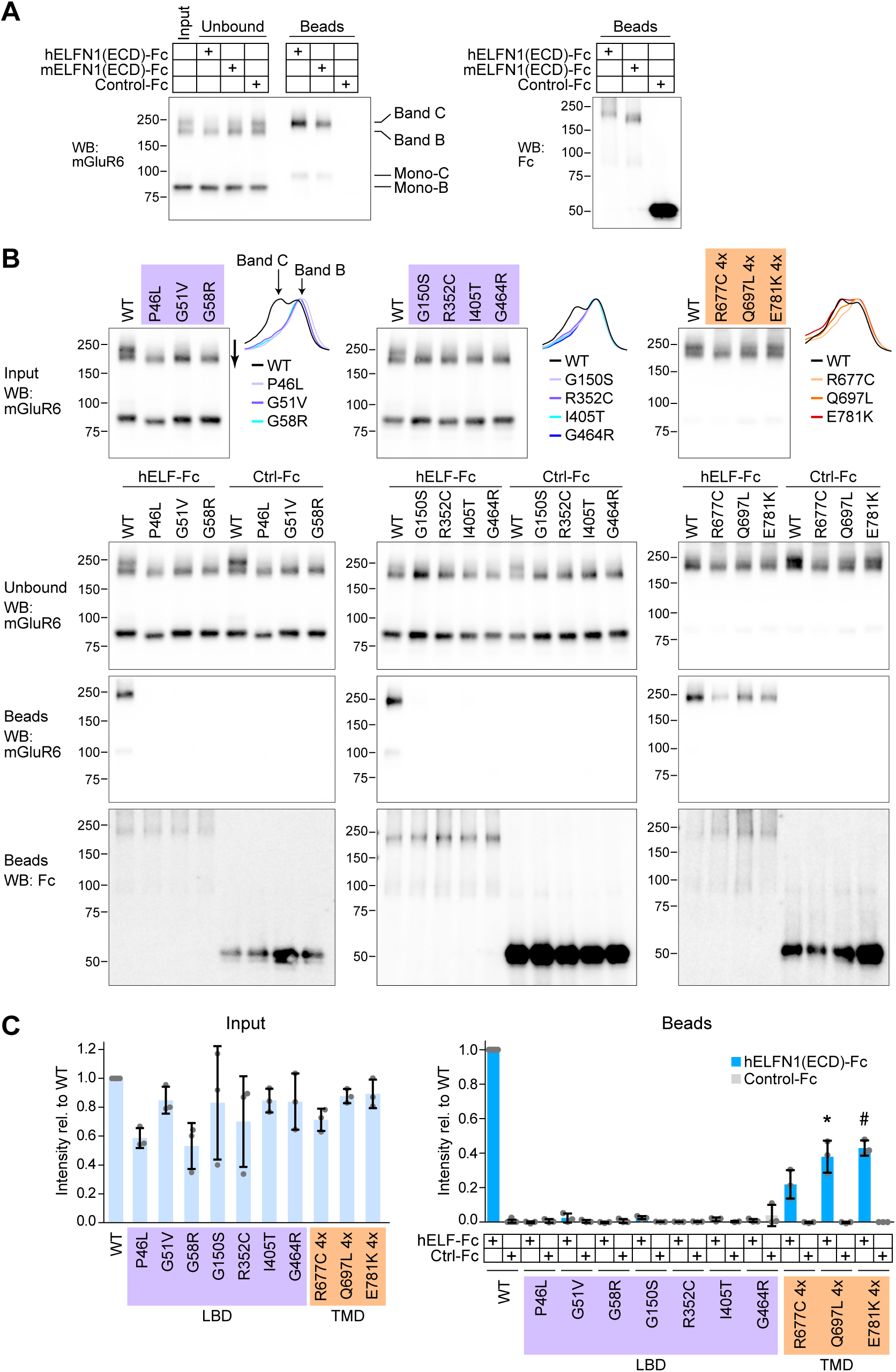
LBD mutants do not bind to ELFN1. **(A)** Pull-down assay with human and mouse ELFN1 extracellular domain (ECD) and human WT mGluR6. ELFN1 ECD fused to Fc, or negative control Fc, was bound to Protein G beads, then mixed with lysate from HEK cells transfected with mGluR6. Samples were subjected to SDS-PAGE and blotted with mGluR6 mAb 1438 or anti-human to detect Fc proteins. Material loaded in bead lanes is ∼5 times the equivalent of material loaded in input and unbound lanes. **(B)** Pull-down assays as in (A), with human ELFN1 ECD and WT or mutant mGluR6. Line profiles of input samples, drawn as shown by the arrow next to the G58R input lane, are shown at right. Profiles are shown normalized to their own minimum and maximum values to highlight differences in width. For TM mutants, four times more transfected cells were used. Westerns were performed with mGluR6 mAb 1438 or mAb 312. **(C)** Quantification of mGluR6 dimer bands in input samples (left) and bead samples (right). Each point represents an independent experiment, and bars represent means ± S.D. ELFN1 bead samples were compared to their corresponding negative control bead samples by two-tailed t test, and p-values were multiplied by 10 to correct for multiple comparisons. *, p<0.05; #, p<0.001.

To confirm that the LBD mutants lack complex glycosylation, HEK cell lysates were treated with Endo H, which removes core N-linked glycans only, or PNGase F, which removes all N-linked glycans. First, the presence of multiple dimer species in control reactions was quantified by measuring the full width at half maximum (FWHM) of line profiles through the dimer bands (Fig 4A-C). The LBD mutants, but not the TMD mutants, had significantly reduced FWHM compared to WT (Fig 4C). As expected, band C in WT samples was Endo H resistant, and this species was lacking in the LBD mutants (Fig 4A). A minor population of Endo H resistant monomer band (Mono-C) was also detected in WT samples only. The amount of Endo H resistant dimer band was quantified by measuring the positive area of the line profile of the Endo H treated sample after subtracting the PNGase F treated sample (Fig 4D,E). All of the tested mutants had significantly less Endo H resistant protein than WT, though the effect was most pronounced for the LBD mutants. ELFN1 binding was correlated with the amount of Endo H resistant protein (Fig 4F), consistent with selective binding to band C.

**Figure 4.**
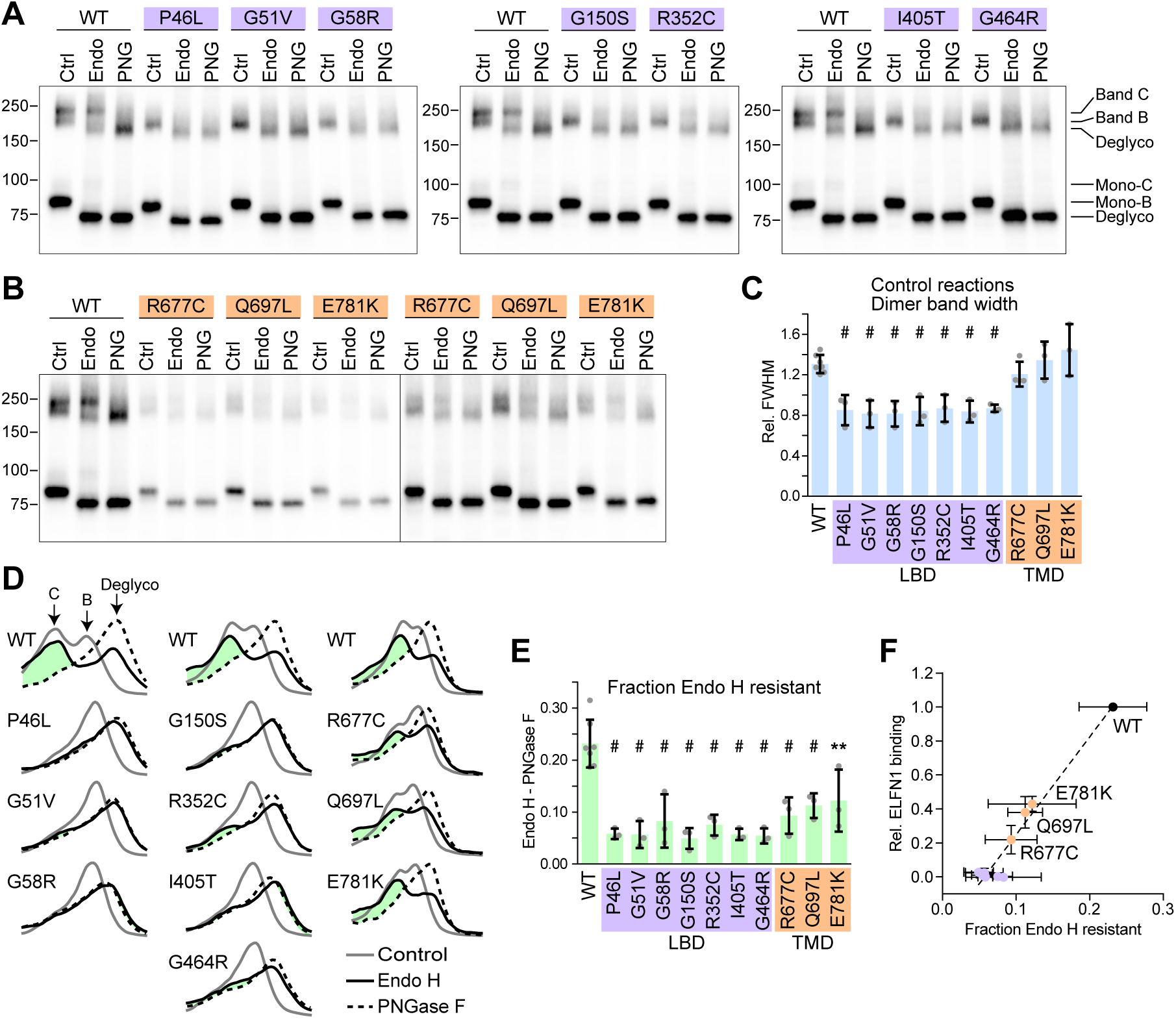
LBD mutants have reduced complex N-glycosylation. **(A)** HEK cell lysates from cells expressing LBD mutants were treated with Endo H or PNGase F, or left untreated (Ctrl), and subjected to SDS-PAGE and blotting with mAb 1438. Positions of Endo H-resistant Band C, Endo H Band B, and the deglycosylated form, as well as their monomer counterparts, are indicated. **(B)** TM mutants were analyzed as in (A). The TM mutant lanes are duplicated on the right side to more clearly show the dimer bands. **(C)** Quantification of full width at half max (FWHM) of line profiles of control lanes in blots such as those shown in (A,B). To factor out variability in migration due to gel running time, values were normalized to the distance between the two peaks of the line profile from the Endo H-treated WT lane on the same blot. **(D)** Line profiles from control, Endo H-treated, and PNGase F-treated lanes from each mGluR6 variant are shown superimposed, after normalizing each profile to the total area under the curve. The positive area remaining after subtracting the PNGase F profile from the Endo H profile is shown in green. **(E)** Quantification of the difference between the Endo H and PNGase F profiles (green areas shown in (D). In (C) and (D), each point represents an independent experiment, and bars represent means ± S.D. Mutants were compared to WT by one-way ANOVA and Dunnett’s post-test. **, p<0.01; #, p<0.001. **(F)** Comparison of the fraction of Endo H-resistant protein (from (E)) and relative ELFN1 binding (from Fig. 3C).

To confirm that decreases in complex glycosylation observed above were not due to prior defects in transfer of the donor glycan to Asn residues in the ER, we measured the Endo H-sensitive monomer band shift after PNGase F treatment (Fig S5). We previously reported that mGluR6 contains four N-glycosylation sites, and when single sites were mutated, the reduction in N-glycan content could be detected by a ∼20-25% reduction in the band shift with PNGase F (17). With the exception of G150S, significant differences in band shifts after PNGase F treatment were not observed here (Fig S5). For G150S, the mean shift was only reduced by ∼13%, and given the similar phenotype of G150S and the other LBD mutants, we conclude that all four N-glycosylation sites are likely occupied.

### LBD mutants employ a Golgi bypass route to the PM

We previously showed that for WT mouse mGluR6, band C, but not the core-glycosylated dimer band B, is present at the PM when expressed in HEK cells (17). However, the robust surface expression of some CSNB mutants lacking complex glycosylation suggests the presence of immature core-glycosylated species at the PM. To test this, we labeled surface proteins with biotin and performed streptavidin bead pull-downs (Fig 5). The dimer bands of WT human mGluR6, similar to mouse mGluR6, were present at the PM primarily in complex-glycosylated form; band C was present in the bead fraction and depleted from the flow-through, while band B was present only in the flow-through. The bead fraction of LBD mutants, in contrast, did contain band B, confirming the presence of core-glycosylated dimer species at the PM. Core-glycosylated monomer bands (Mono-B) were also observed at the PM, for both WT and mutants, and the minor species of slower migrating Endo H-resistant monomer (Mono-C), was also observed for WT. The same samples were blotted for endogenous PM protein Na^+^/K^+^ ATPase as a positive control for surface protein labeling, and α-tubulin as a negative control to demonstrate that intracellular proteins were not labeled. Reactions with the biotinylation reagent omitted did not have any mGluR6 protein in the bead fraction, ruling out nonpecific binding of mGluR6 proteins to the beads.

**Figure 5.**
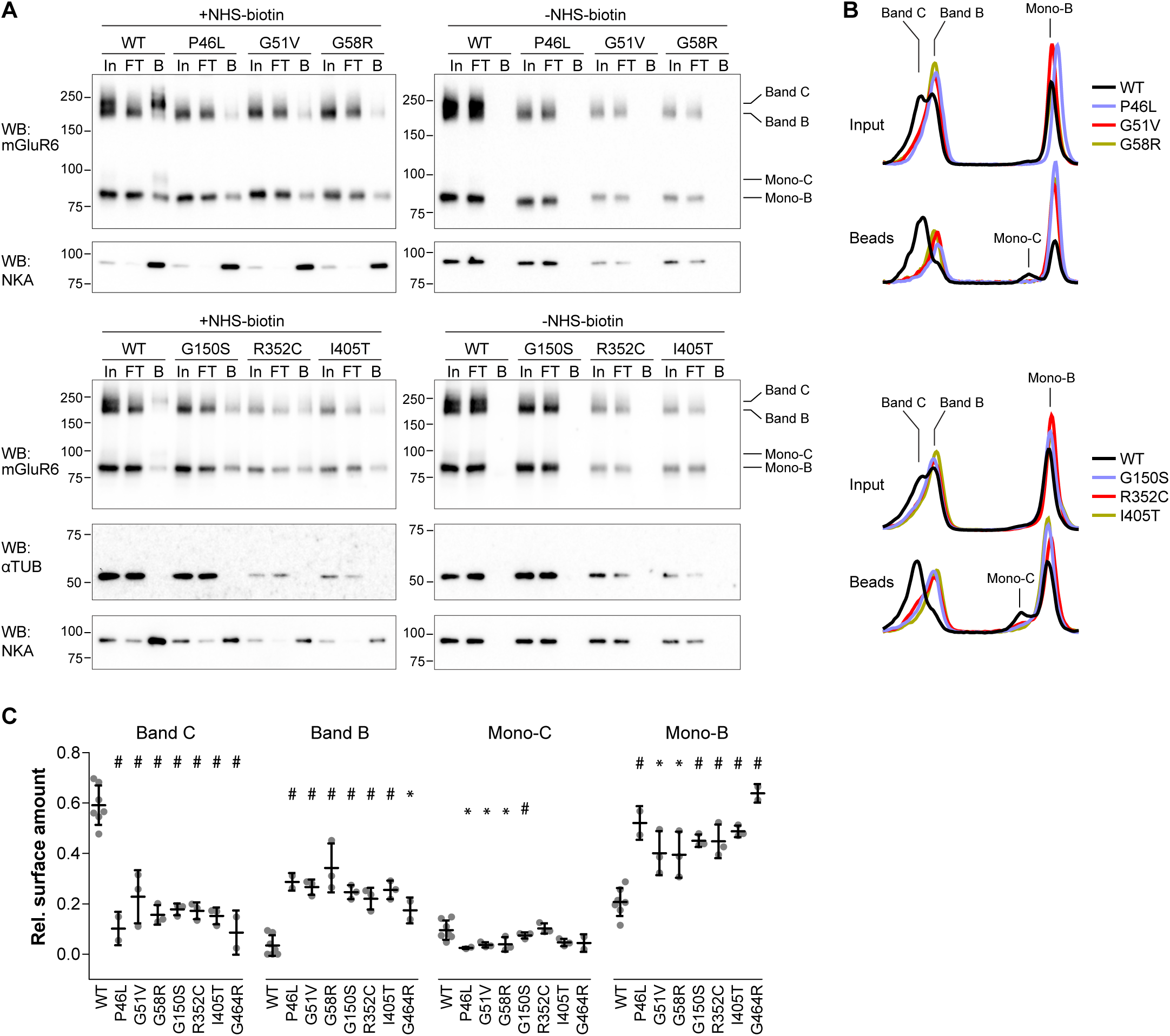
LBD mutants display increased relative amounts of core-glycosylated protein at the PM. **(A)** Live cells were labeled with membrane impermeant biotinylation reagent, followed by quenching and pull-down with streptavidin beads. Input (In), unbound (FT), and 5x equivalent bead (B) samples were resolved by SDS-PAGE and blotted with mAb 1438. The same samples were also blotted for endogenous positive control Na/K ATPase (NKA) and negative control α-Tubulin. **(B)** Example line profiles drawn through bead lanes, including dimer and monomer bands, and shown normalized to total area under the curve. **(C)** Quantification of relative amounts of Band C, Band B, Mono-C, and Mono-B, in bead samples, derived from fitting a sum of four Gaussians to profiles as shown in (B). Each point represents an independent experiment, and bars represent means ± S.D. For each band, mutants were compared to WT by one-way ANOVA and Dunnett’s post-test. *, p<0.05; #, p<0.001.

To quantify the relative amounts of the different mGluR6 species at the PM, line profiles through the bead lanes were normalized to total area under the curve, as shown in Fig 5B, then fit to a sum of four Gaussian distributions (Fig S6). To report the relative amounts of the four species (band C, band B, mono-C, and mono-B), the area under each Gaussian in the fit was calculated (Fig 5C). For all of the LBD mutants, the complement of mGluR6 PM proteins contained greatly reduced amounts of band C, novel appearance of significant amounts of the core glycosylated band B, and increased relative amounts of core-glycosylated monomer. These results demonstrate that trafficking of the mGluR6 LBD mutants is altered such that almost all of the mGluR6 molecules present at the PM are immature core-glycosylated forms.

Maturation of N-linked glycans to Endo H-resistant forms occurs in the Golgi (18, 19). The biotinylation results suggest a trafficking defect whereby the CSNB LBD mutants bypass the Golgi, using a UPS pathway. However, an alternative possibility is that the LBD mutations hinder interaction with Golgi-resident glycosylatransferases. To test this, we treated transfected cells with Brefeldin A (BFA), which leads to fusion of early Golgi compartments with the ER and redistribution of Golgi enzymes into the ER (34–37). For both WT and mutant mGluR6, BFA treatment resulted in a new Endo H resistant dimer band that was similar in apparent size to the Endo H sensitive Band B, as well as a new EndoH resistant monomer band (Fig 6, A-E, Fig S7). The migration of band B in control conditions, as well as the migration of the PNGase F-treated protein, was not affected (Fig 6B). BFA treatment also had no effect on the migration of Endo H resistant WT Band C, with or without Endo H treatment (Fig 6B). These results indicate that the mutants are adequate substrates for at least some Golgi glycosyltransferase enzymes. However, for both WT and mutant mGluR6, the new Endo H resistant product resulting from BFA treatment was not as slow-migrating as band C (Fig 6C), suggesting that additional modification occurs in the trans-Golgi / trans-Golgi network. Furthermore, the new Endo H resistant species of neither WT nor mutant mGluR6 bound efficiently to ELFN1 (Fig 6F), indicating that BFA treatment does not result in the authentic post-translational modification required for ELFN1 binding.

**Figure 6.**
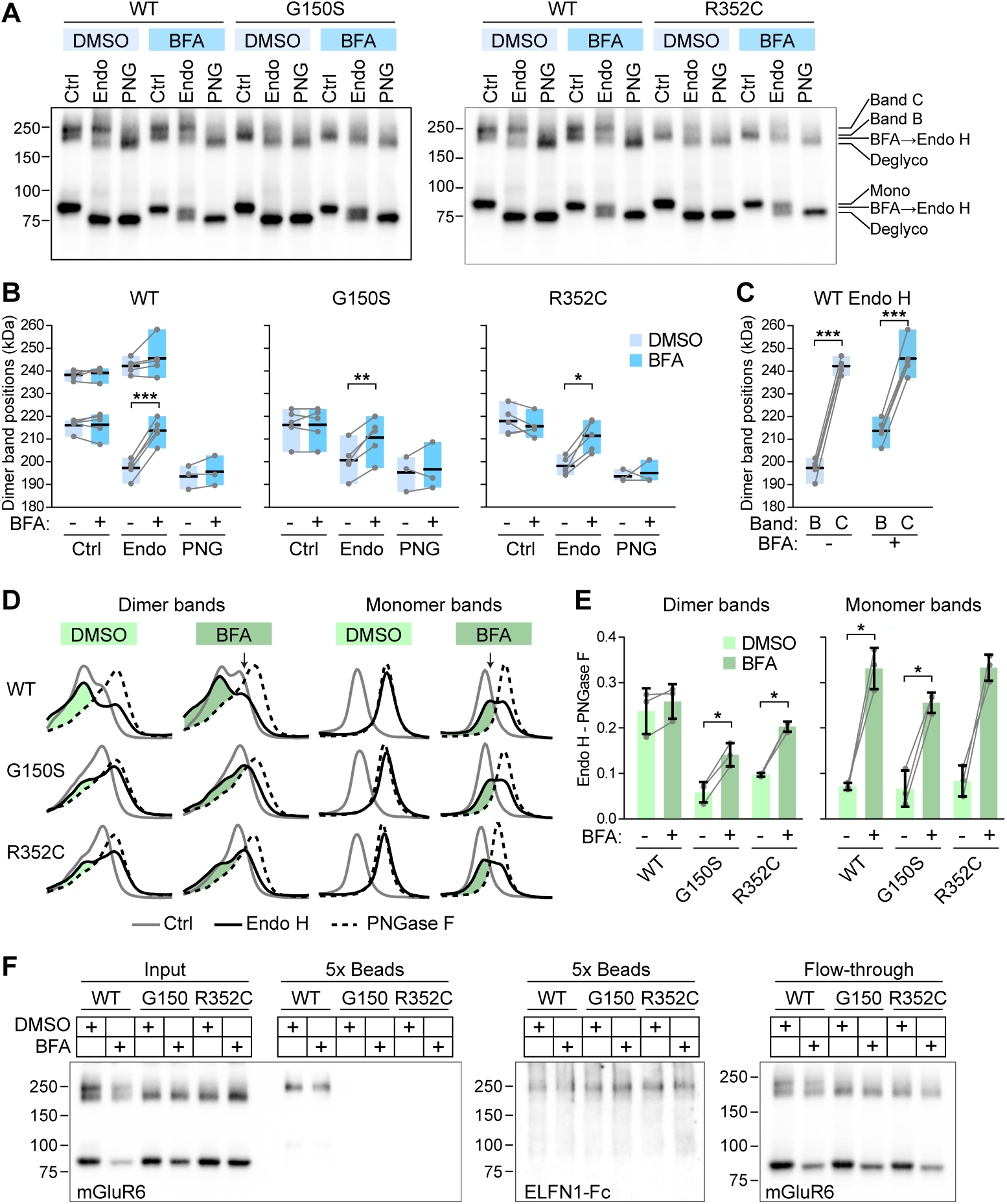
BFA treatment results in new Endo H-resistant band. **(A)** HEK cells were treated with 5 or 20 μg/ml BFA or 0.1% DMSO (vehicle) for 4 h, followed by glycosidase treatment and SDS-PAGE. **(B)** The apparent molecular weight of dimer bands was estimated from peak positions of the line profile and a line fit between flanking marker bands. Peak positions with and without BFA were compared by paired t-test. **(C)** WT Endo H data from (B) is reproduced to compare positions of upper and lower dimer bands by paired t-test. In (B) and (C), bars span minimum to maximum values, points represent independent experiments, and black lines show the means. **(D)** Line profiles through dimer and monomer bands from control, Endo H-treated, and PNGase F-treated lanes from each sample are shown superimposed, after normalizing each profile to the total area under the curve. The positive area remaining after subtracting the PNGase F profile from the Endo H profile is shown in green. **(E)** Quantification of the difference between the Endo H and PNGase F profiles (green areas shown in (D). Endo H resistant protein with and without BFA was compared by paired t-test. Points represent independent experiments, and bars show means ± S.D. P-values from paired t-tests in (B), (C), and (E) were multiplied by three to correct for multiple comparisons. (F) Cells were treated with BFA for 4 h prior to ELFN1 pull-down.

The mechanisms that sort cargo to conventional or UPS pathways are poorly understood. However, ER stress has been shown to upregulate UPS pathways in general (20), and overexpression of mutant proteins could lead to ER stress. To test for this possibility, we blotted lysates from transfected cells for C/EBP-homologous protein (CHOP) (Fig S8), which is upregulated in ER stress conditions (38). No differences in CHOP levels were detected in cells transfected with empty vector, WT mGluR6, or mGluR6 mutants, indicating that overexpression of CSNB mutants likely does not lead to large increases in ER stress. This is consistent with the similar appearance of intracellular WT and mutant mGluR6 in IF experiments (Fig 1), and suggests that the CSNB mutations affect Golgi sorting of mGluR6.

## Discussion

In this study, we report that seven of nine tested CSNB missense mutations in the mGluR6 LBD lead to a glycosylation defect and failure of ELFN1 binding, despite competence for plasma membrane trafficking. Since it is likely that ELFN1 binding is required for synaptic enrichment in BCs (14, 15), this phenotype explains the observed mislocalization of the mutants in BCs, and the electronegative ERG of the CSNB patients. The other two LBD mutations, as well as CRD and TMD mutations, lead to severe reductions in plasma membrane trafficking.

Our results suggest that the glycosylation defect stems from altered sorting in the ER lumen and use of a UPS pathway bypassing the Golgi, where complex N-linked glycans are normally acquired. Results in this study, as well as our previous work (17), are consistent with the requirement for mGluR6 Golgi transit for acquisition of ELFN1 binding ability. The BFA experiments further suggest that the required modification is performed in the trans-Golgi. However, the specific nature of the modification that is required for ELFN1 binding is unknown; it could be an aspect of the complex N-glycans themselves, or a different post-translational modification acquired in concert with glycan maturation.

No structures of mGluR6 have yet been reported. However, the seven LBD mutations leading to Golgi bypass, when mapped to homologous positions in mGluR4 and mGluR7, appear to be located throughout the upper lobe of the LBD (Fig S9). Their disparate locations throughout the lobe suggest they are more likely to cause a loss-of-function for an interaction mediating Golgi sorting, rather than a gain-of-function for an interaction leading to Golgi bypass. Regulation of sorting to the Golgi versus sorting to a UPS pathway is poorly understood. Our results suggest an important role for the upper lobe of the LBD in mGluR6 sorting. On the other hand, there is evidence that upregulation of UPS is mediated by ER stress (20). Though increased ER stress was not detected with the LBD mutants, this possibility cannot be ruled out. Another alternative possibility is that the LBD mutations affect the accessibility of N-glycans to Golgi enzymes for processing. The BFA experiment demonstrated that some complex glycosylation of the LBD mutants can occur, but did not distinguish between glycosylation sites. Interestingly, N451, homologous to mouse N445, which was the N-glycosylated residue that had the greatest loss of band C and defect in ELFN1 binding when mutated (17), is also located in the upper lobe of the LBD (Fig S9). It is possible that the CSNB LBD mutations cause an altered conformation or misfolding of the upper lobe that hinders appropriate modification at N451.

It was previously reported that some CSNB mutants, including P46L, G58R, G150S, and I405T, had no detectable PM protein in HEK cells (24). In contrast, we found that all of these had at least ∼35% of normal PM protein levels, and that of the the G150S mutant in particular was comparable to WT. Our results were highly reproducible, and we obtained similar results with both untagged mGluR6 and mGluR6-EGFP. The source of the discrepancy is unknown, but one possible cause is the use of an N-terminal tag, which could affect trafficking, whereas our assay was performed with mAb 1438, which recognizes an endogenous extracellular epitope, allowing the use of constructs with an authentic N-terminus.

In BCs, the ER extends throughout the soma and dendrites (39). Little is known about secretory trafficking pathways, including the location of Golgi outposts or satellites that might be present in dendrites. We previously showed that endogenous mGluR6 in BCs is mostly complex glycosylated and migrates in SDS-PAGE with a similar apparent size to the upper band in HEK cells (band C) (17). The location of mGluR6 ER exit and Golgi transit in BCs is unknown. While generally immunofluorescence microscopy shows mGluR6 protein restricted to dendritic tips, in certain labeling conditions, mAb 1438 detected endogenous mGluR6 in BC bodies, suggesting that complex glycosylated protein might be present in post-Golgi vesicles throughout the cell body, and possibly the dendrites (32). However, in conditions where WT mGluR6 appears primarily in dendritic tips, the CSNB LBD mutants were mislocalized throughout the cell body and dendrites. The localization pattern is similar to previously reported N-terminal mGluR6 deletion mutants, which were found to be primarily intracellular and partially colocalized with the ER in somas, though colocalization in dendrites was not clear (32). The subcellular location of the CSNB mutants in BCs remains to be determined.

The trafficking and glycosylation defects of CSNB LBD mutants were identified here in heterologous cells. Testing these results in BCs is an important future direction, as is characterizating of the BC dendritic secretory trafficking system in general. Because the low efficiency of plasmid electroporation in BCs precludes detecting the exogenous protein by western blotting, recombinant adeno-associated viruses expressing mGluR6 mutants will be required for biochemical characterization of CSNB mutants expressed in retina. Determining where WT mGluR6 is inserted in the PM, and whether the CSNB mutants can also reach the PM, will also be important for identifying trafficking defects associated with CSNB mutants in BCs.

In summary, we demonstrated that multiple CSNB mutations located in the LBD of mGluR6 confer a similar phenotype, in which proteins reach the plasma membrane lacking the normal complex N-glycosylation, and consequently fail to interact with ELFN1. These mutants are also mislocalized in BCs, consistent with a requirement for trans-synaptic interactions with ELFN1 to mediate synaptic enrichment. The other group III mGluRs (mGluR4, 7, and 8) are also able to interact trans-synaptically with ELFN proteins, and these interactions have roles in synapse formation and function (40–44). All but one of the mutated residues used here are conserved across the group III mGluRs. Studying the effects of homologous mutations in other mGluRs may provide insights into trafficking and ELFN1 interactions, with implications for synapse function throughout the central nervous system.

## Methods

### DNA constructs

Human mGluR6 (NP_000834.2) was PCR amplified from GRM6-Tango (45) (a gift from Bryan Roth; Addgene plasmid # 66391). To make the mGluR6-EGFP fusion, human mGluR6 was fused to a GGGSGGG linker sequence followed by EGFP using overlap extension PCR, and cloned into pCDNA3.1 (for HEK cell expression) or pGrm6P (for BC expression). pGrm6 contains the *Grm6* promoter 200 bp critical region, restricting expression to ON-BCs (46), and was derived from from Addgene plasmid #18817 (a gift from Connie Cepko) as described (39). CSNB mutations were introduced in pGrm6P mGluR6-EGFP by site-directed mutagenesis, and mutant constructs were subsequently cloned in pCDNA3.1, both in untagged and EGFP-fused versions. Untagged human mGluR6 was used unless indicated otherwise. To make N-terminally Myc-tagged human mGluR6, the signal peptide cleavage site was predicted with SignalP-6.0 (47), and the Myc-tag sequence was inserted after the predicted signal sequence, between a.a. 29 and 30, using overlap extension PCR. pCDNA3.1 with untagged mouse mGluR6 (NP_775548.2) was described previously (32, 48). Human ELFN1 cDNA was obtained from the DNASU Plasmid Repository (49) (clone HsCD00861210), and the extracellular domain (a.a. 1-418) was fused to human Fc derived from the BamHI-XbaI fragment of Addgene plasmid #59313 (50) (a gift from Peter Scheiffele and Tito Serafini) with an additional intervening GGAAAA linker. The similarly constructed homologous mouse ELFN1 extracellular domain construct and negative control HAss-flag-Fc constructs were previously described (17).

### Cell culture and transfections

HEK293T cells were maintained without antibiotics in Dulbecco’s modified Eagle medium (DMEM) with 10% fetal bovine serum (Corning) at 37 °C with 5% CO_2_. Cells were seeded in 24-well plates (with poly-D-lysine coated coverslips for IF experiments) and transfected the next day with 0.6 ug DNA and 1 μl Lipofectamine 2000 (Invitrogen) according to the manufacturer’s instructions. For transfection with Fc-tagged proteins, media was changed to DMEM containing 10% ultra-low IgG fetal bovine serum (Avantor Seradigm) before transfection.

### ELFN1 binding assays

Pull-down assays were performed with mouse or human ELFN1 extracellular domain, fused to Fc, as indicated. The procedure was as described previously (17). Media from cells transfected with ELFN1 or negative control Fc was harvested 3 days post-transfection, centrifuged to remove cells, supplemented with 1/20 volumes of 1 M Tris pH 7.4 and a dash of PMSF, and incubated with protein G Plus agarose beads (Calbiochem) with end-over-end mixing for 2 hr at room temperature, then washed with PBS. During the bead binding, cell lysates were prepared from cells transfected with WT or mutant mGluR6 2 days earlier. Cells were washed in PBS and resuspended in ice cold lysis buffer (PBS supplemented with 50 mM NaCl, 1% Triton X-100, and 1× Complete EDTA-free protease inhibitor cocktail (Roche), incubated on ice for 15 min, and centrifuged at 11200 × g for 10 min at 4 °C. An aliquot of supernatant was reserved as the input sample, and the remainder was divided among the ELFN1-Fc and negative control Fc beads. Binding reactions were incubated with end-over-end mixing for 90 min at 4 °C, then washed four times with Wash buffer (PBS supplemented with 50 mM NaCl, 1% Triton X-100, and a dash of PMSF).

### Surface biotinylation

Surface proteins were labeled two days post-transfection. Cells from two wells in a 24-well plate were pooled, washed with PBS-8 (PBS adjusted to pH 8), and incubated for 30 min with with 10 mM EZlink-Sulfo-NHS-SS-biotin (Thermo) dissolved immediately before use in PBS-8, or mock-treated with PBS-8 only. Reactions were quenched with five volumes of 100 mM glycine in PBS-8, then cells were washed twice more in quenching solution, lysed in 50 mM Tris pH 7.4, 200 mM NaCl, 1% Triton X-100, with 1× Complete EDTA-free protease inhibitors (Roche), incubated on ice for 15 min, and centrifuged at 11200×g for 10 min at 4 °C. Supernatants were incubated with streptavidin agarose beads (Thermo) with end-over-end mixing for 90 min at 4 °C, then washed four times with Wash buffer.

### Glycosidase treatments

Cell lysates, prepared as above for pull-down experiments, were treated with 10× glycoprotein denaturing buffer (New England Biolabs) at room temperature for 10 min, then 30 μl denatured lysate were treated with 2 μl Endo H or 1 μl PNGase F enzymes (New England Biolabs) in 40 μl reactions according to the manufacturer instructions, for 1 hr at 37°C. Control reactions were diluted with PBS to 40 μl and incubated on ice.

### Subretinal injection and electroporation

Protocols were approved by the Dalhousie University Committee on Laboratory Animals, and all procedures were performed in accordance with regulations established by the Canadian Council on Animal Care. CD1 albino mice (CD1-Elite #482 were from Charles River Laboratories. Grm6^nob3^ mice (51) (Jackson Laboratory #016883) back-crossed to CD1 (Charles River #022) were previously described (32). For some experiments, nob3 mice were used after one additional back-cross to CD1-Elite. DNA was prepared using a Qiafilter maxiprep kit (Qiagen), and dissolved in water. For injection, DNA solution contained 1× PBS, 0.1% Fast Green FCF dye (Fisher Scientific), 2 mg/ml pGrm6P-mGluR6-EGFP plasmid, and 1 mg/ml pGrm6P DsRed plasmid. Injection and electroporation procedures were essentially as described (33, 52). Briefly, P0 mouse pups were anesthetized on ice, the eyelid was opened, a pilot hole was made with a 30G needle, and and approximately 450 nl of DNA solution was injected into the subretinal space using a 33G blunt needle connected to a UMP3 microinjection syringe pump and MICRO2T controller (World Precision Instruments). Five 50-ms pulses of 80V were applied across the eyes at 1-s intervals using tweezer electrodes and a square wave electroporator (ECM830, BTX Harvard Apparatus). Injected eyes were processed 4-6 weeks later.

### Immunofluorescence microscopy

For labeling HEK293T cells on coverslips, cells were fixed in 2% paraformaldehyde in PBS for 10 min, washed in PBS, then blocked in either PBSA (non-permeabilizing, PBS with 1% BSA) or PBSAT (permeabilizing, PBS with 1% BSA and 0.1% Triton X-100). Cells were labeled with 3 μg/ml mGluR6 mAb 1438 or 3 μg/ml Myc 9E10 diluted in PBSA or PBSAT for 1 h, washed in PBS, then labeled with 2 μg/ml Alexa555-conjugated anti-mouse secondary antibody (Invitrogen), diluted in BPSA or PBSAT, for 30 min, washed in PBS, and mounted with Prolong Diamond (Invitrogen). Images were acquired with a Zeiss LSM880 confocal microscope with 63× oil immersion objective. Alexa555 and EGFP (when present) were imaged sequentially using 561 nm and 488 nm lasers, and 4-6 images were acquired for each coverslip.

Injected eyes were fixed in 2% paraformaldehyde in PBS for ∼20 min, then washed extensively in PBS. Following cryoprotection in 30% sucrose in PBS overnight at 4 °C, corneas were removed and eyecups with lenses were embedded in O.C.T. Compound, and 16 μm sections were collect on Superfrost Plus slides. Sections were rinsed in PBS and blocked for at least 2 hrs in PBS with 10% normal donkey serum, 5% BSA, and 0.2% Triton X-100. Sections were labeled overnight at 4 °C with 8 μg/ml TRPM1 mAb 545H5 diluted in blocking buffer, followed by 8 μg/ml Alexa647-conjugated anti-mouse secondary antibody (Invitrogen) diluted in blocking buffer, for 2 h at room temp. Slides were mounted with Prolong Diamond and z-stacks were acquired (ten images, 0.5 μm z-interval, 1024 × 512 pixels, x-y resolution 65.9 nm/pixel) with a Zeiss LSM880 confocal microscope and 63× oil immersion objective.

### Image analysis

For images of HEK cells, Mathematica v13 (Wolfram) was used to set pixel values ≤ threshold values to zero, as an approximate background correction, using threshold values of 0.03 and 0.01 in the Alexa555 and EGFP channels, respectively, then determine the mean intensity of each image. All values were divided by the average of WT images from the same experiment. OPL puncta in images of retina sections were quantified as previously described (17). Briefly, masked images were made by manually removing the inner nuclear layer, and ROIs corresponding to OPL puncta were derived from scaled and background-corrected masked images using MorphologicalComponents with method Convex Hull in Mathematica. Components containing ≥150 pixels were removed, and resulting ROIs were applied to unscaled images to obtain the intensity within OPL puncta, which was then divided by the total intensity of the unscaled unmasked image.

### CHOP detection

HEK cells were transfected with 0.6 μg empty vector, or WT or mutant mGluR6. As a positive control, untransfected cells were treated with 1 μM thapsigargin (Invitrogen) 4 h prior to harvesting. Cells were pelleted, wash in PBS, and resuspended in 500 μl PBS with 1x Complete EDTA-free protease inhibitor mix (Roche) and 1% phosphatase inhibitor cocktail 3 (Sigma). After addition of ¼ volume of 5× sample buffer, samples were passed ∼20 times through a 26G needle and subjected to SDS-PAGE.

### Western blotting

Samples were mixed with ¼ volume of 5× sample buffer (250 mM Tris pH 6.8, 10% SDS, 50% glycerol, 10% β-mercaptoethanol, bromophenol blue dye) and loaded without heating onto Tris-glycine SDS-PAGE gels. Proteins were transferred to nitrocellulose, and membranes were blocked in 5% milk in TBST (50 mM Tris-HCl, 150 mM NaCl, 0.1% Tween-20, pH 8.4), incubated with primary antibodies at 4 °C overnight, washed in TBST, incubated with secondary antibody (horseradish peroxidase conjugated goat anti-mouse (Jackson Immunoresearch, 0.08 or 0.16 μg/ml), horseradish peroxidase conjugated goat anti-rabbit (Proteintech, 0.02 or 0.04 μg/ml), for 2 hrs at room temperature, and washed in TBST. Fc fusion proteins were detected with DyLight 680 conjugated goat anti-human (Thermo #SA510138, 0.25 μg/ml) incubated overnight as a primary antibody. SuperSignal West Pico PLUS Chemiluminescent Substrate (Thermo) was added, and blots were imaged with an Azure 500 digital imager with 2x2 binning.

### Western blot analysis

For ELFN1 binding experiments, band intensities were estimated by measuring the integrated density of a box drawn around the band, subtracted by a background box of the same size placed either above or below the band. For analysis of band positions and relative amounts of mGluR6 species, line profiles from 10-px wide lines drawn through each lane in ImageJ were analyzed. Profiles were normalized to the sum of all points in the curve (∼ area under the curve). Apparent molecular weights of bands were estimated using the peak positions and a linear fit between the peak positions of two flanking marker bands. Peak positions were identified using the FindPeaks function in Mathematica. In cases where one of the dimer doublet peaks appeared as a shoulder, peak positions were manually estimated. All peak positions used for calculation of apparent molecular weights in Fig 6 are shown in Fig S7. For analysis of relative amounts of mGluR6 species, line profiles were first normalized to the total area under the curve. For estimation of amounts of mGluR6 species present in bead samples following biotinylation experiments (Fig 5), area-normalized line profiles including both dimer and monomer bands were fit to a sum of four Gaussian distributions 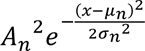 with constraints 0 < *σ*_*n*_ < 10 pixels and each *μ*_*n*_ within 1 px of peak positions identified from the ensemble of bead samples on one gel, including WT. Relative amounts of band C, band B, mono-C, and mono-B species were determined by numerical integration of each Gaussian distribution.

### Primary antibodies

mGluR6 mAbs developed with mouse mGluR6 were previously described and validated for IF and westerns using nob3 retina tissue (32) and transfected HEK cells (17). Comparison of the mGluR6 mAbs for detection of mouse and human mGluR6 is shown in Fig S1. mGluR6 mAbs were used at 1 μg/ml for western blotting and 3 μg/ml for IF. TRPM1 mAb 545H5 was previously described and validated with Trpm1^-/-^ retina tissue (53, 54). Purification of mGluR6 and TRPM1 antibodies from hybridoma cell culture media using protein G Sepharose Fast Flow (Cytiva) was previously described (32). For later antibody preparations, hybridomas were first adapted to low-serum media, and purified using 90% Cell mAb Medium (Gibco), 8.5% Iscove’s Modified Dulbecco’s medium, and 1.5% ultralow IgG serum (Avantor Seradigm) (mAbs 312, 1438, and 545H5), or 75% Cell mAb Medium, 21.3% Iscove’s, and 3.8% ultralow IgG serum (mAb 366), along with 100 U/ml penicillin and 100 μg/ml streptomycin. Purification of Myc tag clone 9E10 from hybridomas obtained from the Developmental Studies Hybridoma Bank (The University of Iowa) was previously described (32). The following commercial antibodies were used: Na/K ATPase (ATPA1 mAb M7-PB-E9, Invitrogen #MA3-928, 1-2 μg/ml), α-Tubulin (mAb DM1A, Cell Signaling #3873, 1:1000), CHOP (Proteintech #15204-1-AP, 0.7 μg/ml).

## Acknowledgements

This work was supported by an Alcon Young Investigator Grant (to M.A.A.), Natural Sciences and Engineering Research Council of Canada (NSERC) Discovery Grant RGPIN-2022-02982 (to M.A.A.) and the Dalhousie Department of Ophthalmology & Visual Sciences and Dalhousie Medical Research Foundation (DMRF) New Faculty Research Grant (to M.A.A.) funded by donations to DMRF including a contribution from Novartis. The work was also supported by the Canada Foundation for Innovation John R. Evans Leaders Fund and Research Nova Scotia. Students were partially supported by NSERC Undergraduate Summer Research Awards (M.L.M. and M.P.), Dalhousie Department of Physiology & Biophysics studentships (J.L. and A.P.R.), and a Dr. R. Evatt and Rita Mathers Trainee Scholarship (F.A.K.A.).

## Supplementary Figure Legends

**Fig S1. Comparison of mGluR6 mAb detection of mouse (Ms) and human (Hu) mGluR6 in HEK cells. (A)** Cells transiently transfected with mouse or human mGluR6, or empty vector, were labeled in permeabilizing or non-permeabilizing conditions with mGluR6 mAbs as indicated. Images of non-permeabilized samples were acquired and processed with different settings. **(B)** Lysates from cells transfected with mouse or human mGluR6 were blotted with mGluR6 mAbs or tubulin antibody. **(C)** Sequence alignmnent of human mGluR6 with previously identified epitopes in mouse mGluR6 (32). Non-identical residues are shown in red.

**Fig S2. Surface expression of human mGluR6-EGFP mutants in HEK cells. (A)** Transfected cells were labeled in non-permeabilizing conditions with mAb 1438 to detect surface protein. EGFP fluorescence was imaged to detect total protein. For poorly expressed mutants, the EGFP images are reproduced rescaled (bottom row) to demonstrate the presence of transfected cells. **(B)** Quantification of surface and total protein in transfected HEK cells. Values were normalized to the mean of WT samples from the same day, and points represent means of at least three images each from independent experiments; bars show means +/- S.D. Mutants were compared to the WT value of 1 using 1-sample t-tests, and p-values were multiplied by 16 to correct for multiple comparisons. *, p<0.05; **, p<0.01.

**Fig S3. Co-transfection of mutant mGluR6 had no effect on surface expression of WT ss-Myc-mGluR6. (A,B)** Cells were co-transfected with 0.4 μg WT ss-Myc-mGluR6 and 0.4 μg either empty vector (EV) or indicated untagged WT or mutant mGluR6. Example images are shown. **(C)** Quantification of images as shown in (A-B). Each point is the mean of four images from an independent experiment. Error bars show means ± S.E.M.

**Fig S4. LBD mutants are mislocalized in nob3 mice. (A)** Example images of retinas electroporated with mGluR6-EGFP and labeled with TRPM1 antibody. **(B)** Magnified views of boxed regions in (A). **(C)** Diagram of electroporation procedure. **(D-F)** Quantification of images as shown in (A). Data are shown relative to the mean of nob3 mice electroporated with WT mGluR6-EGFP. Mutants were compared to WT by one-way ANOVA and Dunnett’s post-test; #, p<0.001. **(G)** Comparison of TRPM1 puncta/total intensity in WT CD1 and nob3 mice. Data are shown relative to the mean of WT CD1 mice electroporated with WT mGluR6-EGFP. Right, aggregated TRPM1 puncta/total and TRPM1 total intensity data for all images. WT and nob3 mice were compared by two-tailed t test. In (D-G), each point represents the mean of at least three images from a different animal (n=2-4 biological replicates per condition), and bars represent means of biological replicates ± S.D.

**Fig S5. Monomer band shifts after PNGase treatment.** The distance between peak positions of monomer bands in control and PNGase F lanes are shown relative to WT on the same blot. Lanes from PNGase experiments shown in Figure 4 and DMSO control samples in BFA experiments shown in Figure 6 were combined. Bars show means +/- S.D. Values were compared to 1 using 1-sample t-tests, and p-values were multiplied by 10 to correct for multiple comparisons. *, p<0.05.

**Fig S6. Gaussian fits used in** Figure 5C. Line profiles (black lines) of bead fractions after surface biotinylation experiments were fit to a sum of four Gaussians (red). The sum and four Gaussian terms are shown.

**Fig S7.** Peak positions in line profiles used in Figure 6.

**Fig S8. CHOP protein levels are similar in cells transfected with WT and mutant mGluR6.** Cells were transfected with empty vector (EV), WT mGluR6, or indicated mGluR6 mutants. Untransfected cells treated with ∼1 μM thapsigargin (Tg) were used as a positive control. Whole cell homogenates were subjected to SDS-PAGE and blotted with CHOP and α-tubulin antibodies. **(A)** Example western blots. **(B)** Quantification of CHOP/tubulin, relative to WT. Each point is the mean of duplicate lanes from an independent experiment; error bars show means ± S.E.M.

**Fig S9.** Homologous positions of the seven CSNB Golgi bypass mutations (red) and N-glycosylation sites (cyan) in mGluR4 **(A)** (PDB 7E9H (55)) and mGluR7 **(B)** (PDB 7EPC (56)). N451, which is the homologous position in human mGluR6 to the N445 glycosylation site in mouse mGluR6, is indicated. Extracellular domains are shown.

## References

1. Nakajima Y, Iwakabe H, Akazawa C, Nawa H, Shigemoto R, Mizuno N, Nakanishi S. 1993. Molecular characterization of a novel retinal metabotropic glutamate receptor mGluR6 with a high agonist selectivity for L-2-amino-4-phosphonobutyrate. J Biol Chem 268, 11868–11873.

2. Nomura A, Shigemoto R, Nakamura Y, Okamoto N, Mizuno N, Nakanishi S. 1994. Developmentally regulated postsynaptic localization of a metabotropic glutamate receptor in rat rod bipolar cells. Cell 77, 361–369.

3. Masu M, Iwakabe H, Tagawa Y, Miyoshi T, Yamashita M, Fukuda Y, Sasaki H, Hiroi K, Nakamura Y, Shigemoto R, Takada M, Nakamura K, Nakao K, Katsuki M, Nakanishi S. 1995. Specific deficit of the ON response in visual transmission by targeted disruption of the mGluR6 gene. Cell 80, 757–765.

4. Koyasu T, Kondo M, Miyata K, Ueno S, Miyata T, Nishizawa Y, Terasaki H. 2008. Photopic electroretinograms of mGluR6-deficient mice. Curr Eye Res 33, 91–99.

5. Vardi N. 1998. Alpha Subunit of G(o) Localizes in the Dendritic Tips of ON Bipolar Cells. J Comp Neurol 395, 43–52.

6. Nawy S. 1999. The metabotropic receptor mGluR6 may signal through G(o), but not phosphodiesterase, in retinal bipolar cells. J Neurosci 19, 2938–2944.

7. Dhingra A, Lyubarsky A, Jiang M, Pugh EN, Birnbaumer L, Sterling P, Vardi N. 2000. The light response of ON bipolar neurons requires Gαo. J Neurosci 20, 9053–9058.

8. Morgans CW, Zhang J, Jeffrey BG, Nelson SM, Burke NS, Duvoisin RM, Brown RL. 2009. TRPM1 is required for the depolarizing light response in retinal ON-bipolar cells. Proc Natl Acad Sci U S A 106, 19174–19178.

9. Shen Y, Heimel JA, Kamermans M, Peachey NS, Gregg RG, Nawy S. 2009. A transient receptor potential-like channel mediates synaptic transmission in rod bipolar cells. J Neurosci 29, 6088–6093.

10. Koike C, Obara T, Uriu Y, Numata T, Sanuki R, Miyata K, Koyasu T, Ueno S, Funabiki K, Tani A, Ueda H, Kondo M, Mori Y, Tachibana M, Furukawa T. 2010. TRPM1 is a component of the retinal ON bipolar cell transduction channel in the mGluR6 cascade. Proc Natl Acad Sci USA 107, 332–337.

11. Koike C, Numata T, Ueda H, Mori Y, Furukawa T. 2010. TRPM1: A vertebrate TRP channel responsible for retinal ON bipolar function. Cell Calcium 48, 95–101.

12. Shen Y, Rampino MAF, Carroll RC, Nawy S. 2012. G-protein–mediated inhibition of the Trp channel TRPM1 requires the Gβγ dimer. Proc Natl Acad Sci U S A 109, 8752–8757.

13. Xu Y, Orlandi C, Cao Y, Yang S, Choi C-I, Pagadala V, Birnbaumer L, Martemyanov KA, Vardi N. 2016. The TRPM1 channel in ON-bipolar cells is gated by both the α and the βγ subunits of the G-protein Go. Sci Rep 6, 20940.

14. Cao Y, Sarria I, Fehlhaber KE, Kamasawa N, Orlandi C, James KN, Hazen JL, Gardner MR, Farzan M, Lee A, Baker S, Baldwin K, Sampath AP, Martemyanov KA. 2015. Mechanism for selective synaptic wiring of rod photoreceptors into the retinal circuitry and its role in vision. Neuron 87, 1248–1260.

15. Cao Y, Wang Y, Dunn HA, Orlandi C, Shultz N, Kamasawa N, Fitzpatrick D, Li W, Zeitz C, Hauswirth W, Martemyanov KA. 2020. Interplay between cell-adhesion molecules governs synaptic wiring of cone photoreceptors. Proc Natl Acad Sci U S A 117, 23914–2392.

16. Ellaithy A, Gonzalez-Maeso J, Logothetis DA, Levitz J. 2020. Structural and biophysical mechanisms of class C G protein-coupled receptor function. Trends Biochem Sci 45, 1049–1064.

17. Miller ML, Pindwarawala M, Agosto MA. 2024. Complex N-glycosylation of mGluR6 is required for trans-synaptic interaction with ELFN adhesion proteins. J Biol Chem 300, 107119.

18. Esmail S, Manolson MF. 2021. Advances in understanding N-glycosylation structure, function, and regulation in health and disease. Eur J Cell Biol 100, 151186.

19. Grieve AG, Rabouille C. 2011. Golgi bypass: Skirting around the heart of classical secretion. Cold Spring Harb Perspect Biol 3, a005298.

20. Gee HY, Kim J, Lee MG. 2018. Unconventional secretion of transmembrane proteins. Semin Cell Dev Biol 83, 59–66.

21. Zeitz C, Robson AG, Audo I. 2015. Congenital stationary night blindness: an analysis and update of genotype-phenotype correlations and pathogenic mechanisms. Prog Retin Eye Res 45, 58–110.

22. Zeitz C, van Genderen M, Neidhardt J, Luhmann UFO, Hoeben F, Forster U, Wycisk K, Mátyás G, Hoyng CB, Riemslag F, Meire F, Cremers FPM, Berger W. 2005. Mutations in GRM6 cause autosomal recessive congenital stationary night blindness with a distinctive scotopic 15-Hz flicker electroretinogram. Invest Ophthalmol Vis Sci 46, 4328–4335.

23. Dryja TP, McGee TL, Berson EL, Fishman GA, Sandberg MA, Alexander KR, Derlacki DJ, Rajagopalan AS. 2005. Night blindness and abnormal cone electroretinogram ON responses in patients with mutations in the GRM6 gene encoding mGluR6. Proc Natl Acad Sci U S A 102, 4884–4889.

24. Zeitz C, Forster U, Neidhardt J, Feil S, Kalin S, Leifert D, Flor PJ, Berger W. 2007. Night blindness-associated mutations in the ligand-binding, cysteine-rich, and intracellular domains of the metabotropic glutamate receptor 6 abolish protein trafficking. Hum Mutat 28, 771–780.

25. Zeitz C, Labs S, Lorenz B, Forster U, Uksti J, Kroes HY, de Baere E, Leroy BP, Cremers FPM, Wittmer M, van Genderen MM, Sahel JA, Audo I, Poloschek CM, Mohand-Saïd S, Fleischhauer JC, Hüffmeier U, Moskova-Doumanova V, Levin A V., Hamel CP, Leifert D, Munier FL, Schorderet DF, Zrenner E, Friedburg C, Wissinger B, Kohl S, Berger W. 2009. Genotyping microarray for CSNB-associated genes. Investig Ophthalmol Vis Sci 50, 5919–5926.

26. Sergouniotis PI, Robson AG, Li Z, Devery S, Holder GE, Moore AT, Webster AR. 2012. A phenotypic study of congenital stationary night blindness (CSNB) associated with mutations in the GRM6 gene. Acta Ophthalmol 90, e192--e197.

27. Wang Q, Gao Y, Li S, Guo X, Zhang Q. 2012. Mutation screening of TRPM1, GRM6, NYX and CACNA1F genes in patients with congenital stationary night blindness. Int J Mol Med 30, 521–526.

28. Liu H-Y, Huang J, Xiao H, Zhang M-J, Shi F-F, Jiang Y-H, Du H, He Q, Wang Z-Y. 2019. Pseudodominant inheritance of autosomal recessive congenital stationary night blindness in one family with three co-segregating deleterious GRM6 variants identified by next-generation sequencing. Mol Genet Genomic Med 7, e952.

29. Tourville A, Michiels C, Condroyer C, Meunier A, Cordonnier M, Sahel JA, Audo I, Abramowicz M, Zeitz C. 2019. Identification of a novel GRM6 mutation in a previously described consanguineous family with complete congenital stationary night blindness. Ophthalmic Genet 40, 182–184.

30. Huang L, Bai X, Xie Y, Zhou Y, Wu J, Li N. 2024. Clinical and genetic studies for a cohort of patients with congenital stationary night blindness. Orphanet J Rare Dis 19, 101.

31. Kim AH, Liu PK, Chang YH, Kang EYC, Wang HH, Chen N, Tseng YJ, Seo GH, Lee H, Liu L, Chao AN, Chen KJ, Hwang YS, Wu WC, Lai CC, Tsang SH, Hsiao MC, Wang NK. 2022. Congenital stationary night blindness: Clinical and genetic features. Int J Mol Sci 23, 14965.

32. Agosto MA, Adeosun AA, Kumar N, Wensel TG. 2021. The mGluR6 ligand-binding domain, but not the C-terminal domain, is required for synaptic localization in retinal ON-bipolar cells. J Biol Chem 297, 101418.

33. Agosto MA, Wensel TG. 2021. LRRTM4 is a member of the transsynaptic complex between rod photoreceptors and bipolar cells. J Comp Neurol 529, 221–233.

34. Lippincott-Schwartz J, Yuan LC, Bonifacino JS, Klausner RD. 1989. Rapid redistribution of Golgi proteins into the ER in cells treated with brefeldin A: Evidence for membrane cycling from Golgi to ER. Cell 56, 801–813.

35. Lippincott-Schwartz J, Yuan L, Tipper C, Amherdt M, Orci L, Klausner RD. 1991. Brefeldin A’s effects on endosomes, lysosomes, and the TGN suggest a general mechanism for regulating organelle structure and membrane traffic. Cell 67, 601–616.

36. Chege NW, Pfeffer SR. 1990. Compartmentation of the Golgi complex: Brefeldin-A distinguishes trans-Golgi cisternae from the trans-Golgi network. J Cell Biol 111, 893–899.

37. Ladinsky MS, Howell KE. 1992. The trans-Golgi network can be dissected structurally and functionally from the cisternae of the Golgi coplex by brefeldin A. Eur J Cell Biol 59, 92–105.

38. Chen X, Shi C, He M, Xiong S, Xia X. 2023. Endoplasmic reticulum stress: molecular mechanism and therapeutic targets. Signal Transduct Target Ther 8, 352.

39. Agosto MA, Anastassov IA, Robichaux MA, Wensel TG. 2018. A large endoplasmic reticulum-resident pool of TRPM1 in retinal ON bipolar cells. eNeuro 5, e0143.

40. Tomioka NH, Yasuda H, Miyamoto H, Hatayama M, Morimura N, Matsumoto Y, Suzuki T, Odagawa M, Odaka YS, Iwayama Y, Won Um J, Ko J, Inoue Y, Kaneko S, Hirose S, Yamada K, Yoshikawa T, Yamakawa K, Aruga J. 2014. Elfn1 recruits presynaptic mGluR7 in trans and its loss results in seizures. Nat Commun 5, 1–16.

41. Dunn HA, Patil DN, Cao Y, Orlandi C, Martemyanov KA. 2018. Synaptic adhesion protein ELFN1 is a selective allosteric modulator of group III metabotropic glutamate receptors in trans. Proc Natl Acad Sci USA 115, 5022–5027.

42. Dunn HA, Zucca S, Dao M, Orlandi C, Martemyanov KA. 2019. ELFN2 is a postsynaptic cell adhesion molecule with essential roles in controlling group III mGluRs in the brain and neuropsychiatric behavior. Mol Psychiatry 24, 1902–1919.

43. Stachniak TJ, Sylwestrak EL, Scheiffele P, Hall BJ, Ghosh A. 2019. Elfn1-induced constitutive activation of mGluR7 determines frequency-dependent recruitment of somatostatin interneurons. J Neurosci 39, 4461–4474.

44. Park D, Park S, Song J, Kang M, Lee S, Horak M, Suh YH. 2020. N-linked glycosylation of the mGlu7 receptor regulates the forward trafficking and transsynaptic interaction with Elfn1. FASEB J 34, 14977–14996.

45. Kroeze WK, Sassano MF, Huang XP, Lansu K, McCorvy JD, Giguère PM, Sciaky N, Roth BL. 2015. PRESTO-Tango as an open-source resource for interrogation of the druggable human GPCRome. Nat Struct Mol Biol 22, 362–369.

46. Kim DS, Matsuda T, Cepko CL. 2008. A core paired-type and POU homeodomain-containing transcription factor program drives retinal bipolar cell gene expression. J Neurosci 28, 7748–7764.

47. Teufel F, Almagro Armenteros JJ, Johansen AR, Gíslason MH, Pihl SI, Tsirigos KD, Winther O, Brunak S, von Heijne G, Nielsen H. 2022. SignalP 6.0 predicts all five types of signal peptides using protein language models. Nat Biotechnol 40, 1023–1025.

48. Kang HJ, Menlove K, Ma J, Wilkins A, Lichtarge O, Wensel TG. 2014. Selectivity and evolutionary divergence of metabotropic glutamate receptors for endogenous ligands and G proteins coupled to phospholipase C or TRP channels. J Biol Chem 289, 29961–29974.

49. Seiler CY, Park JG, Sharma A, Hunter P, Surapaneni P, Sedillo C, Field J, Algar R, Price A, Steel J, Throop A, Fiacco M, Labaer J. 2014. DNASU plasmid and PSI:Biology-Materials repositories: Resources to accelerate biological research. Nucleic Acids Res 42, D1253–D1260.

50. Scheiffele P, Fan J, Choih J, Fetter R, Serafini T. 2000. Neuroligin expressed in nonneuronal cells triggers presynaptic development in contacting axons. Cell 101, 657–669.

51. Maddox DM, Vessey KA, Yarbrough GL, Invergo BM, Cantrell DR, Inayat S, Balannik V, Hicks WL, Hawes NL, Byers S, Smith RS, Hurd R, Howell D, Gregg RG, Chang B, Naggert JK, Troy JB, Pinto LH, Nishina PM, McCall MA. 2008. Allelic variance between GRM6 mutants, Grm6nob3 and Grm6nob4 results in differences in retinal ganglion cell visual responses. J Physiol 586, 4409–4424.

52. Matsuda T, Cepko CL. 2004. Electroporation and RNA interference in the rodent retina in vivo and in vitro. Proc Natl Acad Sci U S A 101, 16–22.

53. Agosto MA, Anastassov IA, Wensel TG. 2018. Differential epitope masking reveals synapse-specific complexes of TRPM1. Vis Neurosci 35, e001.

54. Agosto MA, Zhang Z, He F, Anastassov IA, Wright SJ, McGehee J, Wensel TG. 2014. Oligomeric state of purified transient receptor potential melastatin-1 (TRPM1), a protein essential for dim light vision. J Biol Chem 289, 27019–27033.

55. Lin S, Han S, Cai X, Tan Q, Zhou K, Wang D, Wang X, Du J, Yi C, Chu X, Dai A, Zhou Y, Chen Y, Zhou Y, Liu H, Liu J, Yang D, Wang MW, Zhao Q, Wu B. 2021. Structures of Gi-bound metabotropic glutamate receptors mGlu2 and mGlu4. Nature 594, 583–588.

56. Du J, Wang D, Fan H, Xu C, Tai L, Lin S, Han S, Tan Q, Wang X, Xu T, Zhang H, Chu X, Yi C, Liu P, Wang X, Zhou Y, Pin JP, Rondard P, Liu H, Liu J, Sun F, Wu B, Zhao Q. 2021. Structures of human mGlu2 and mGlu7 homo- and heterodimers. Nature 594, 589–593.

