## Supplemental Figures for "Defective glycosylation and ELFN1 binding of mGluR6 congenital stationary night blindness mutants"

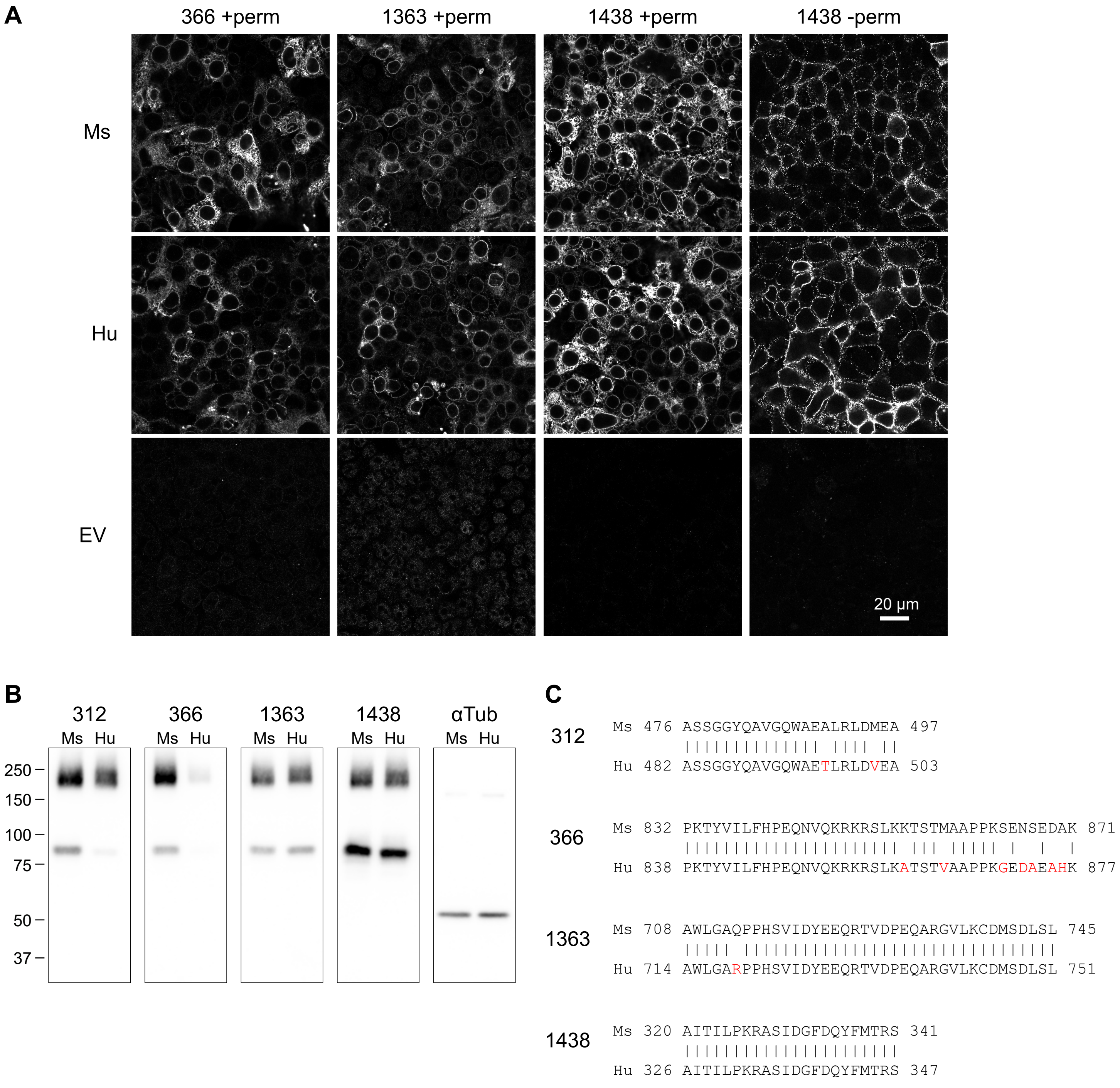


**Fig S1. Comparison of mGluR6 mAb detection of mouse (Ms) and human (Hu) mGluR6 in HEK cells. (A)** Cells transiently transfected with mouse or human mGluR6, or empty vector, were labeled in permeabilizing or non-permeabilizing conditions with mGluR6 mAbs as indicated. Images of non-permeabilized samples were acquired and processed with different settings. **(B)** Lysates from cells transfected with mouse or human mGluR6 were blotted with mGluR6 mAbs or tubulin antibody. **(C)** Sequence alignmnent of human mGluR6 with previously identified epitopes in mouse mGluR6 (Agosto et al. 2021). Non-identical residues are shown in red.

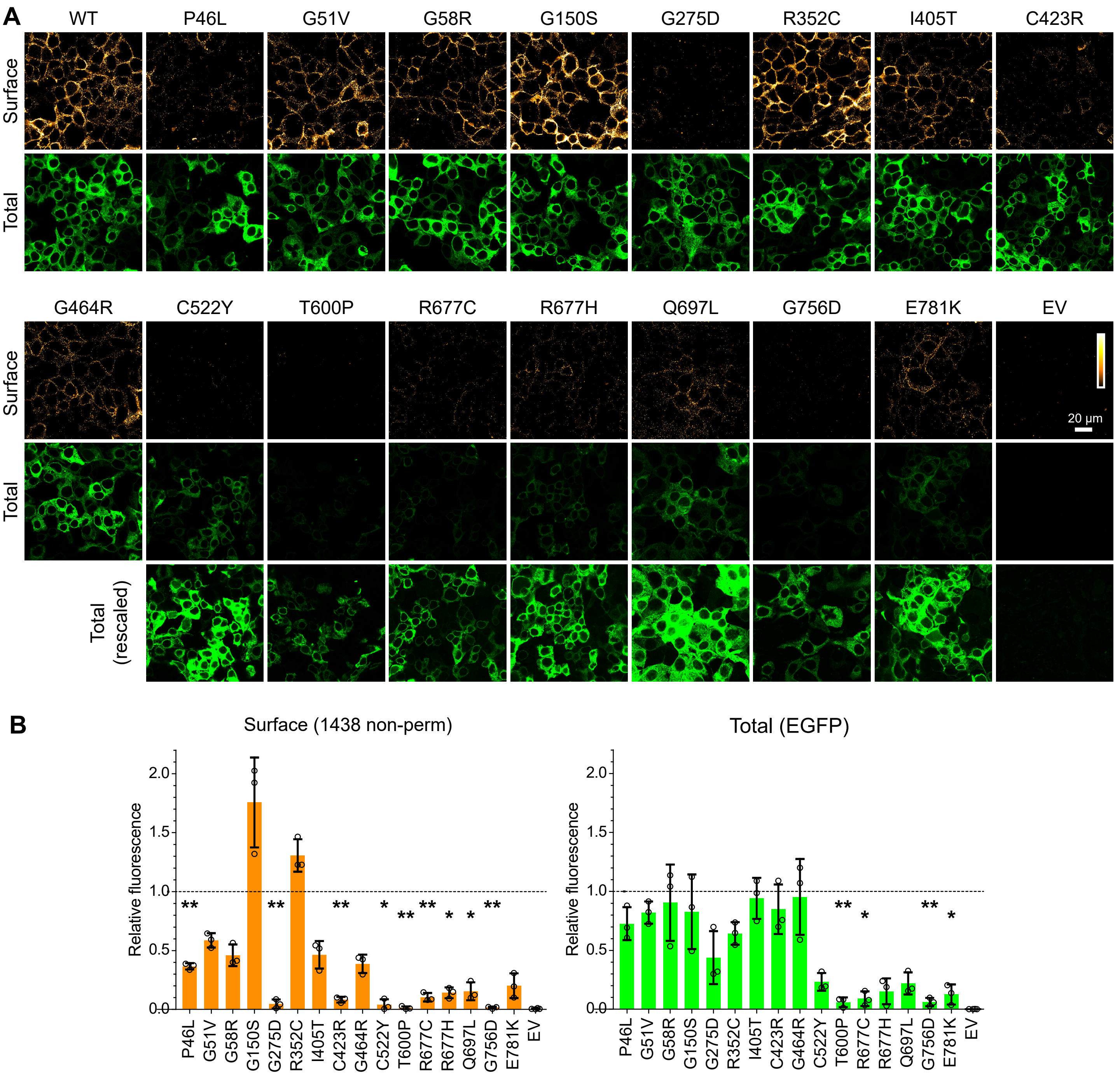


**Fig S2. Surface expression of human mGluR6-EGFP mutants in HEK cells. (A)** Transfected cells were labeled in non-permeabilizing conditions with mAb 1438 to detect surface protein. EGFP fluorescence was imaged to detect total protein. For poorly expressed mutants, the EGFP images are reproduced rescaled (bottom row) to demonstrate the presence of transfected cells. **(B)** Quantification of surface and total protein in transfected HEK cells. Values were normalized to the mean of WT samples from the same day, and points represent means of at least three images each from independent experiments; bars show means +/- S.D. Mutants were compared to the WT value of 1 using 1-sample t-tests, and p-values were multiplied by 16 to correct for multiple comparisons. *, p<0.05; **, p<0.01.

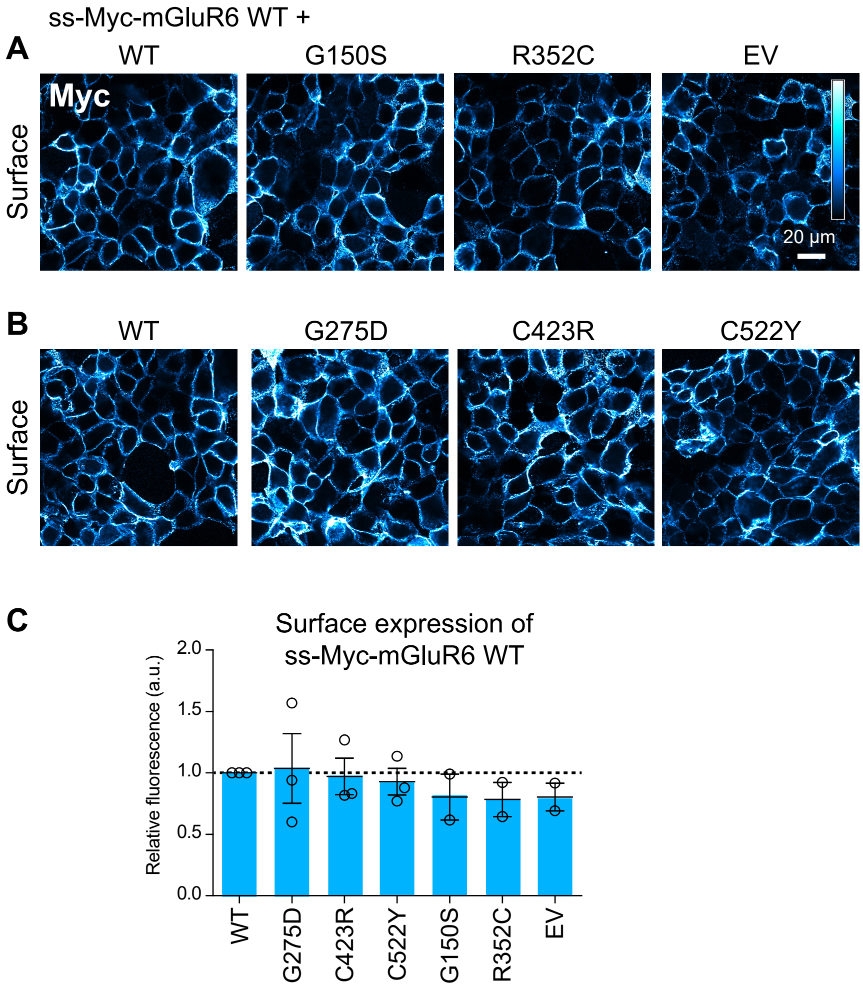


**Fig S3. Co-transfection of mutant mGluR6 had no effect on surface expression of WT ss-Myc-mGluR6. (A,B)** Cells were co-transfected with 0.4 µg WT ss-Myc-mGluR6 and 0.4 µg either empty vector (EV) or indicated untagged WT or mutant mGluR6. Example images are shown. **(C)** Quantification of images as shown in (A-B). Each point is the mean of four images from an independent experiment. Error bars show means ± S.E.M.

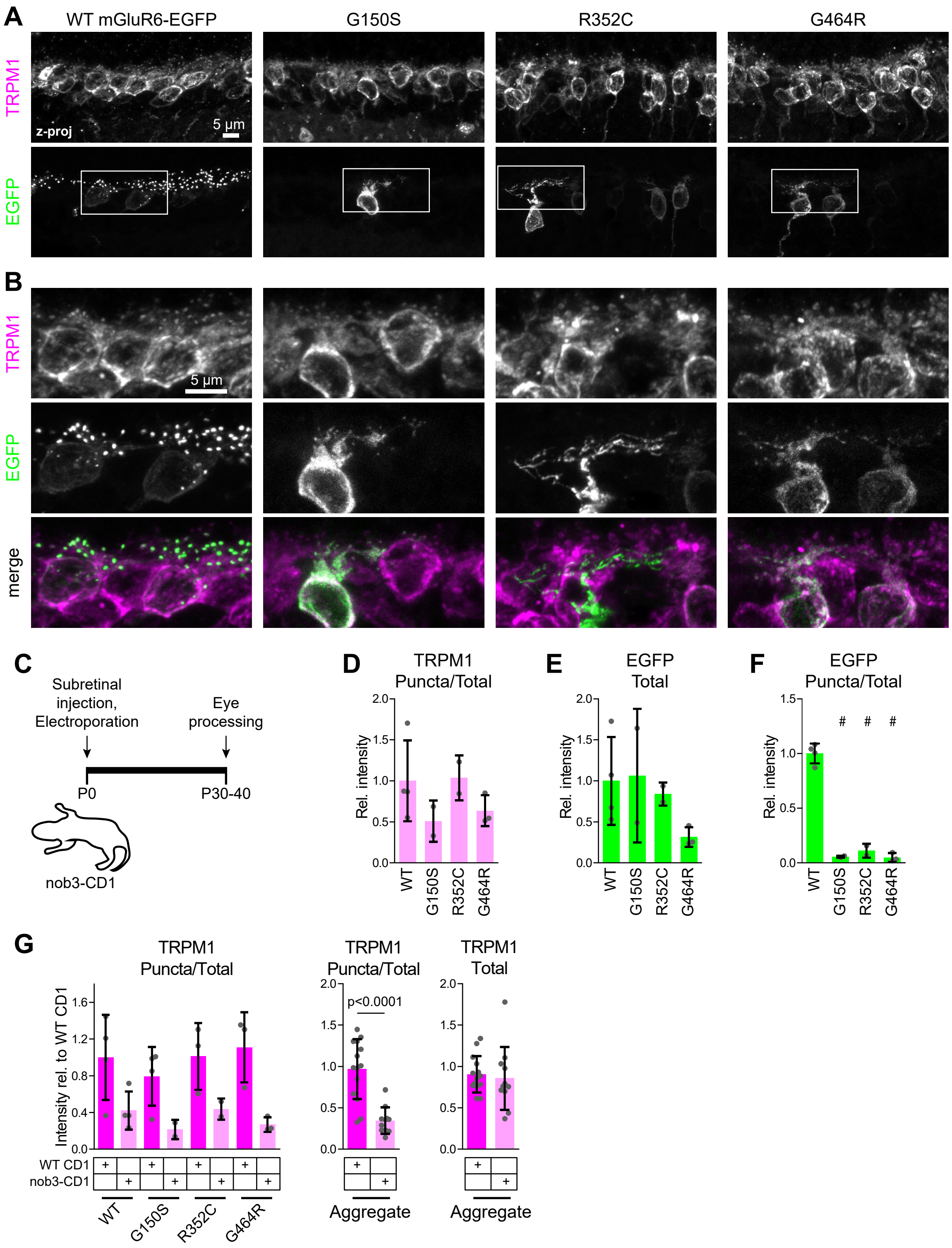


**Fig S4. LBD mutants are mislocalized in nob3 mice. (A)** Example images of retinas electroporated with mGluR6-EGFP and labeled with TRPM1 antibody. **(B)** Magnified views of boxed regions in (A). **(C)** Diagram of electroporation procedure. **(D-F)** Quantification of images as shown in (A). Data are shown relative to the mean of nob3 mice electroporated with WT mGluR6-EGFP. Mutants were compared to WT by one-way ANOVA and Dunnett's post-test; #, $p<0.001$. **(G)** Comparison of TRPM1 puncta/total intensity in WT CD1 and nob3 mice. Data are shown relative to the mean of WT CD1 mice electroporated with WT mGluR6-EGFP. Right, aggregated TRPM1 puncta/total and TRPM1 total intensity data for all images. WT and nob3 mice were compared by two-tailed t test. In (D-G), each point represents the mean of at least three images from a different animal (n=2-4 biological replicates per condition), and bars represent means of biological replicates ± S.D.

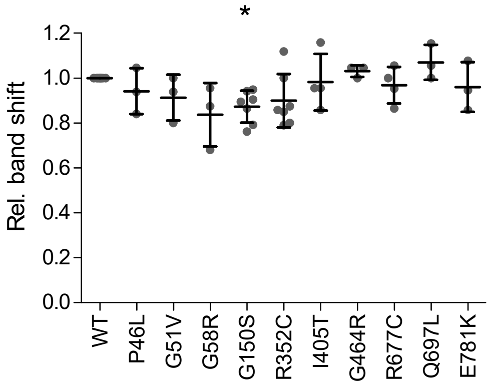


**Fig S5. Monomer band shifts after PNGase treatment.** The distance between peak positions of monomer bands in control and PNGase F lanes are shown relative to WT on the same blot. Lanes from PNGase experiments shown in Figure 4 and DMSO control samples in BFA experiments shown in Figure 6 were combined. Bars show means +/- S.D. Values were compared to 1 using 1-sample t-tests, and p-values were multiplied by 10 to correct for multiple comparisons. *, p<0.05.

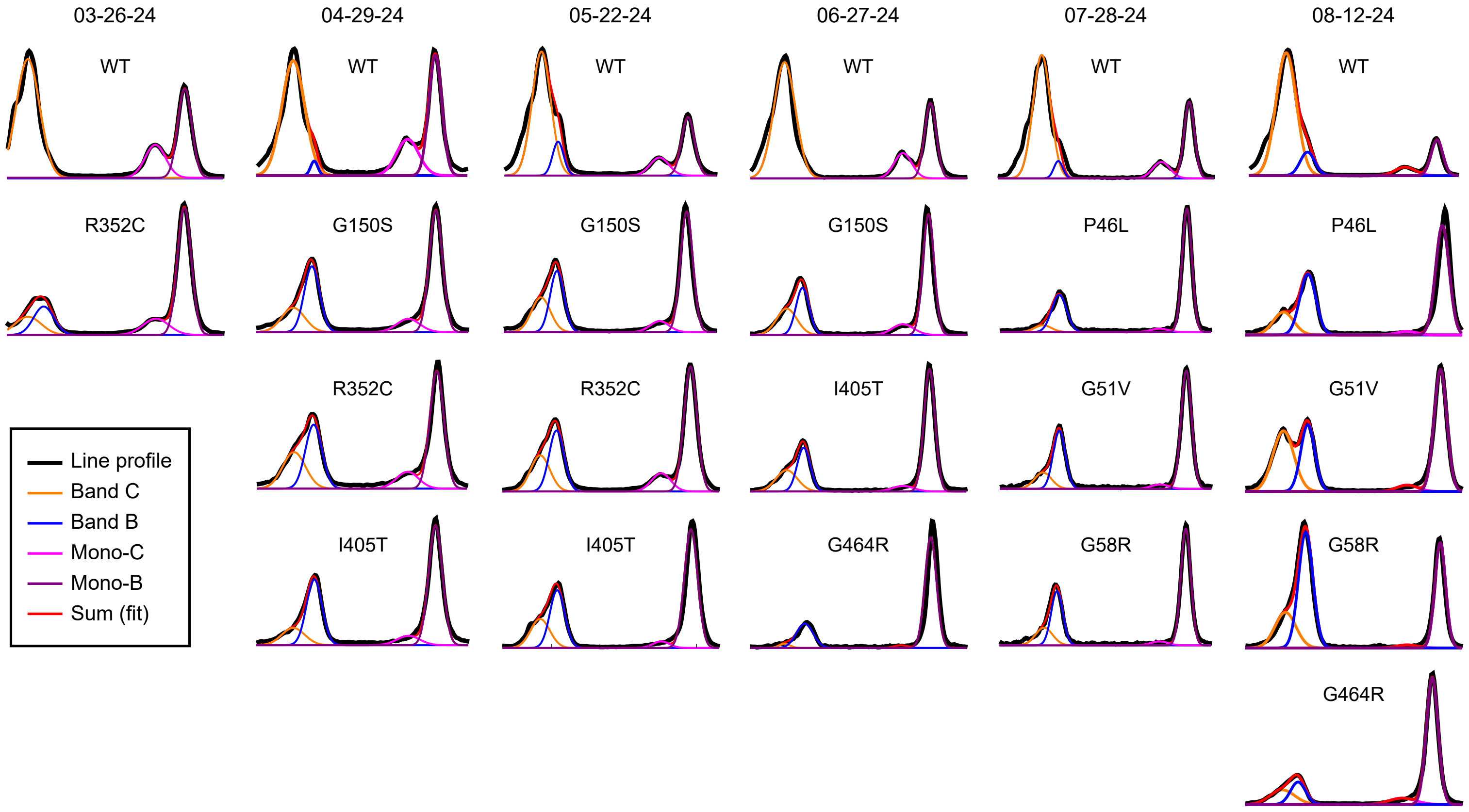


**Fig S6. Gaussian fits used in Figure 5C.** Line profiles (black lines) of bead fractions after surface biotinylation experiments were fit to a sum of four Gaussians (red). The sum and four Gaussian terms are shown.

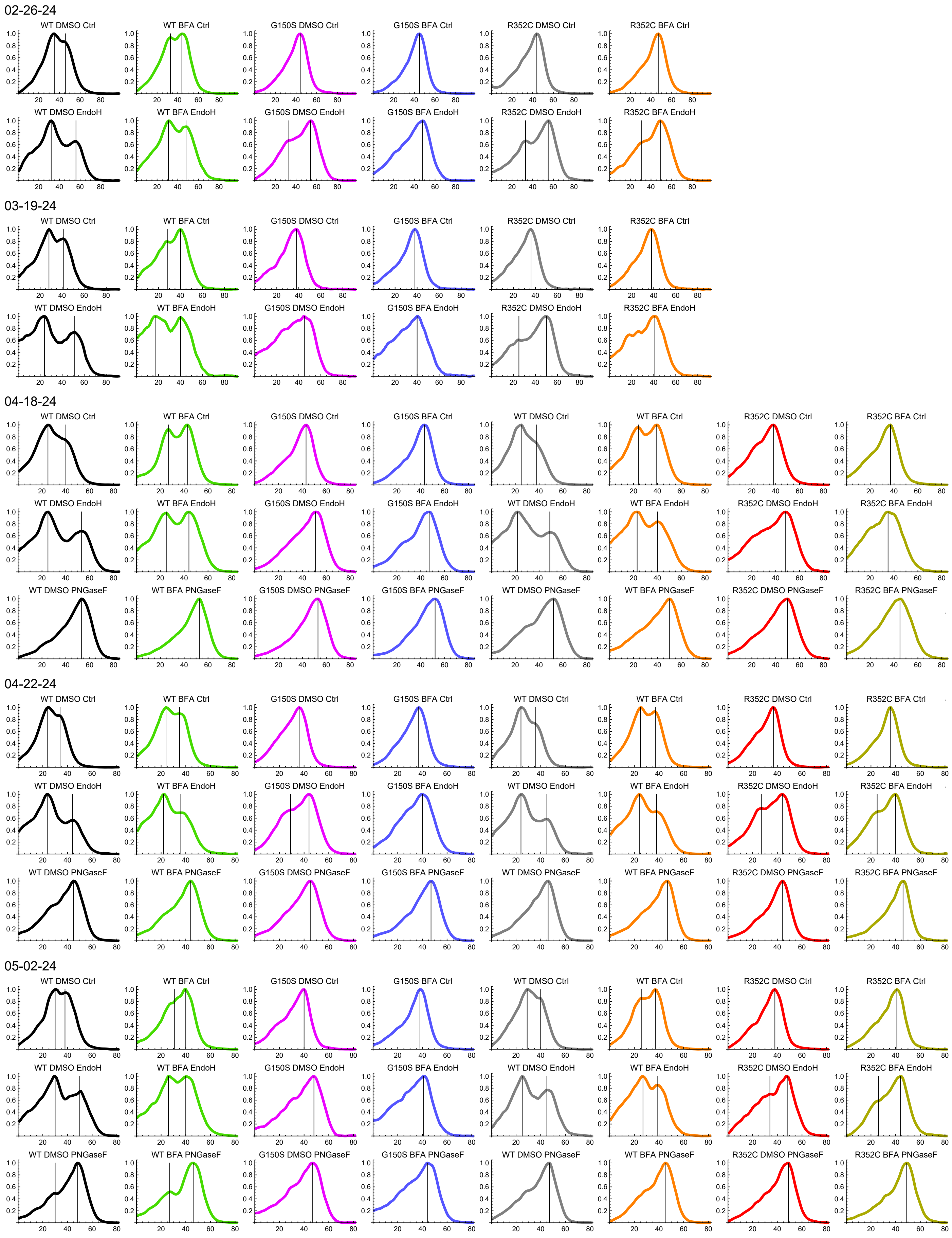


**Fig S7.** Peak positions in line profiles used in Figure 6.

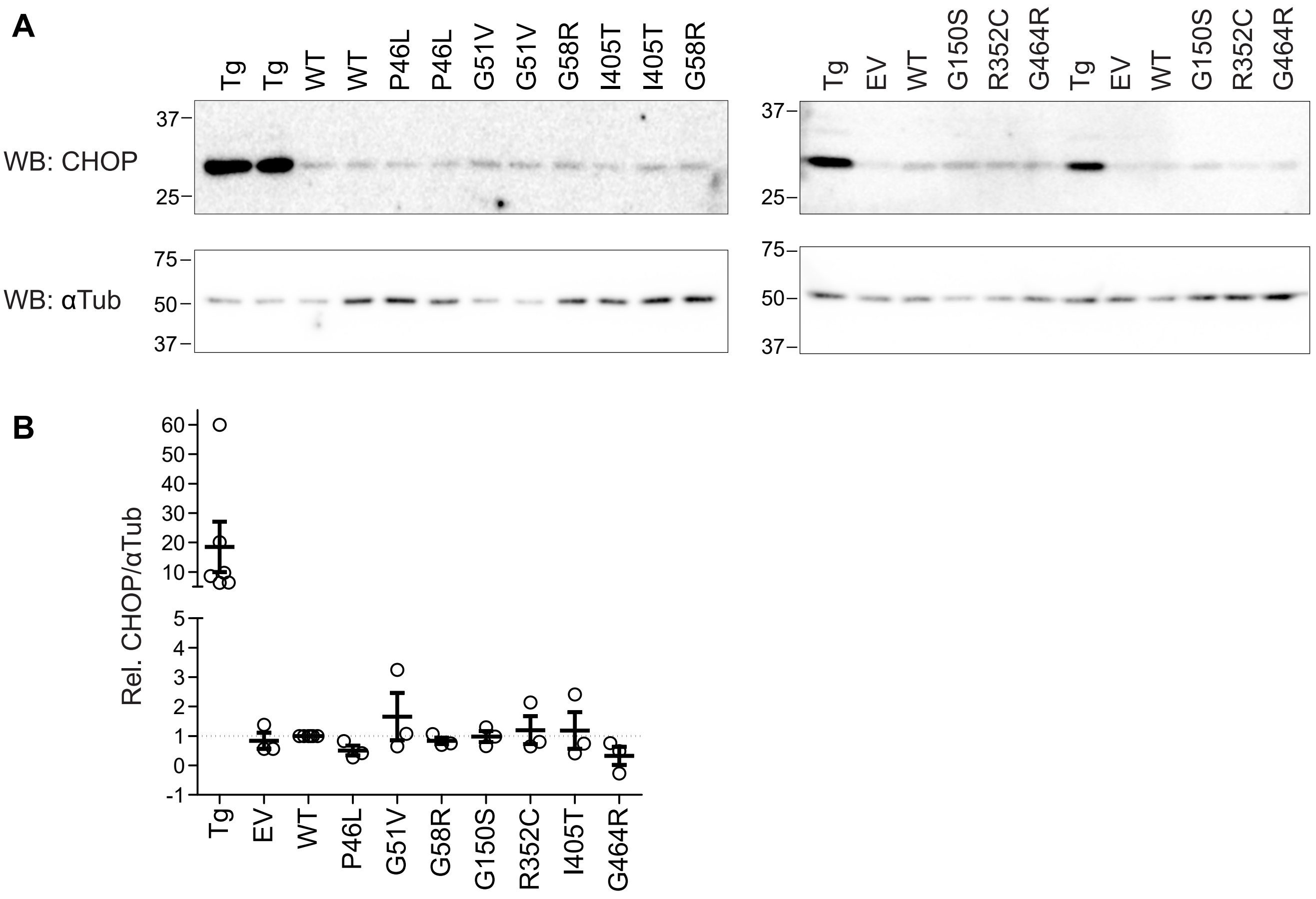


**Fig S8. CHOP protein levels are similar in cells transfected with WT and mutant mGluR6.** Cells were transfected with empty vector (EV), WT mGluR6, or indicated mGluR6 mutants. Untransfected cells treated with ~1 μM thapsigargin (Tg) were used as a positive control. Whole cell homogenates were subjected to SDS-PAGE and blotted with CHOP and α-tubulin antibodies. **(A)** Example western blots. **(B)** Quantification of CHOP/tubulin, relative to WT. Each point is the mean of duplicate lanes from an independent experiment; error bars show means ± S.E.M.

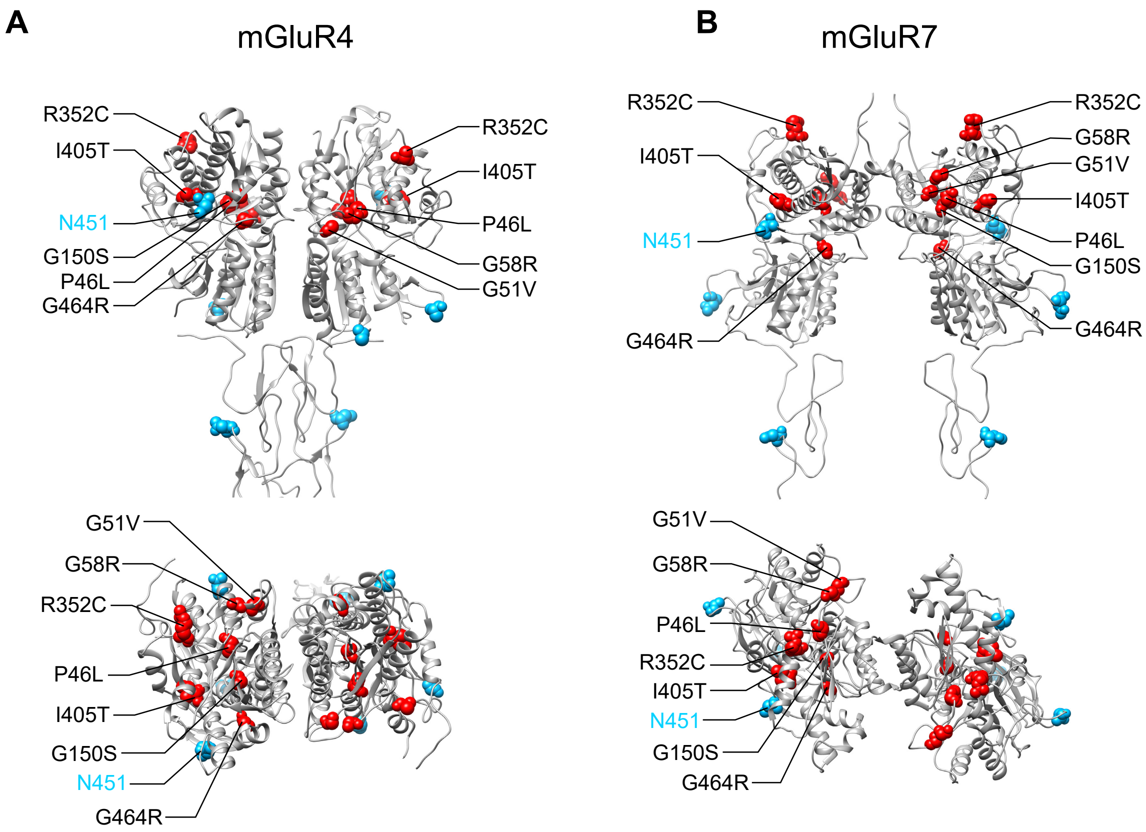


**Fig S9.** Homologous positions of the seven CSNB Golgi bypass mutations (red) and N-glycosylation sites (cyan) in mGluR4 **(A)** (PDB 7E9H, Lin et al. 2021) and mGluR7 **(B)** (PDB 7EPC, Du et al. 2021). N451, which is the homologous position in human mGluR6 to the N445 glycosylation site in mouse mGluR6, is indicated. Extracellular domains are shown.
